# Local macrophage and cDC1 cooperation in the liver drives T-cell immune surveillance against pioneering metastatic cancer cells

**DOI:** 10.64898/2026.11.21.768283

**Authors:** Beatrice Pinci, Almudena Manzanal, María L. Chiclana-Valcárcel, Carlos Luri-Rey, Enric Vercher, David Causapé, Saray Garasa, Mercedes Leal, Leticia Odriozola, Nekane Soria, Arantza Azpilikueta, Elixabet Bolaños, Raluca Alexandru, Karmele Valencia, Ana Rouzaut, María J Garrido, Teresa Lozano, Alfonso Calvo, Carlos E. de Andrea, Mario Squadrito, Sean R. Collins, Pedro Berraondo, Andrés Hidalgo, Ignacio Melero, Alvaro Teijeira

## Abstract

The liver is one of the most frequent sites for hematogenous metastasis across different types of cancer, but how the liver mounts effective immune responses against arriving tumor cells is not fully understood. Using intravital imaging, we have visualized the initial steps of liver metastasis and delineated three phases in the immune response at the earliest stages of liver metastasis: first, there is continuous surveillance of the liver parenchyma by discrete immune cell subsets in the premetastatic tissue. Second, upon metastatic cell arrival, there is massive neutrophil recruitment and tumor-cell material uptake mediated by resident macrophages. Third, antitumor-specific T-cell activation takes place in situ within intrahepatic leukocyte clusters. We find that tumor-derived material uptake triggers transcriptional programs in liver myeloid cells that upregulate chemoattraction programs in Kupffer cells and cDCs, leading to T-cell activation. Antigen cross-presentation and a functional partnership between cDC1s and local macrophages to recruit and form T-cell activation clusters are required for the elicitation of these intrahepatic T-cell responses. Consequently, prior expansion of cDCs by Flt3L-Ig treatment significantly enhances immunosurveillance, curtailing successful metastasis development. Collectively, our findings identify coordinated immune responses that are essential for anti-metastatic immunity within the liver, uncovering a program to mount defenses against tumor cells arriving in the liver.

## INTRODUCTION

The liver is characterized by a structurally compartmentalized resident myeloid population of distinct macrophage subsets and dendritic cells (DCs) deployed to respond to pathogens arriving through different routes (Patseas, et al. 2025). The positioning of myeloid populations is defined by liver morphology, liver vasculature and metabolic zonation of hepatocytes (Gola, et al. 2021; Guilliams, et al. 2022). The liver capsule is covered by macrophages and DCs, which can respond to potential infectious and inflammatory risks in the peritoneal cavity (Sierro, et al. 2017). In the steady state, liver sinusoids are also populated by a specialized population of macrophages with enhanced phagocytic capacities, termed Kupffer cells (KCs), that reside on top of the liver sinusoidal fenestrated endothelial cells (Araujo David, et al. 2026). Within the liver parenchyma, resident DCs are confined to the perivascular space of the vessels present in the periportal triads (Watanabe, et al. 2007). Other minoritarian macrophage populations characterized by their metabolic functions have also been identified within the liver parenchyma (Nusse and Kubes 2025). The specific functions of these myeloid populations have not yet been well delineated across all pathological conditions, but, as in other tissues, antigen-presenting cell functions range from maintenance of tissue homeostasis and elimination of invading pathogens to antigen presentation and immune regulation.

Studies on liver immunosurveillance in the context of infection and inflammation have shown that different local adaptive immune responses can be elicited within the liver by the coordinate activity of distinct non-immune and immune resident cells and by newly recruited leukocytes from the blood. Studies in viral infection have shown that infected hepatocytes can drive direct priming of T cells, resulting in anergic T cell responses (Benechet, et al. 2019; Guidotti, et al. 2015). However, local priming driven by KCs during Hepatitis B (HBV) infection is needed to elicit proper anti-viral cytotoxic T lymphocytes (CTLs), which requires cooperation of CD4 T cells via CD40L (Benechet, et al. 2019; Venzin, et al. 2025). Importantly, local activation of CTLs within the liver results in the formation of specialized activating clusters (termed iMATEs by the researcher who first described them), similar to those found in other tissues, such as the skin (Huang, et al. 2013). Likewise, tertiary lymphoid structure-like immune cell accumulations have been described in HCC, forming around tumor lesions and are important for the outcome of immunotherapy responses (Shu, et al. 2024). Therefore, the liver can be a site for effective generation of adaptive immune responses in adequate inflammatory conditions.

The liver is also one of the most commonly metastasized organs in cancer. Although gastrointestinal cancer liver metastasis is the most frequent, other tumor types, such as lung cancer, melanoma, or breast and bladder cancer, often metastasize to this organ (Tsilimigras, et al. 2021). The two main factors that likely explain this enhanced susceptibility of the liver to metastasis are the sinusoidal and fenestrated structure of liver vasculature (Huu Hoang, et al. 2022) and the immuno-privileged status of the liver that protects it from excessive inflammation caused by the intestinal microbiota and hepatic metabolites (Racanelli and Rehermann 2006).

Liver metastasis is a condition that, when not amenable to surgery, leads to markedly reduced survival of the cancer patient. Importantly, the presence of liver metastasis seems to strongly correlate with a lack of response to immunotherapies, such as immune checkpoint blockade (Lee, et al. 2020; Yu, et al. 2021).

Experimental liver metastasis models have shown the interplay of several leukocyte populations in the immune response elicited by the presence of cancer cells arriving in the liver and implications in the immune surveillance of liver metastasis. In models of liver metastasis using melanoma cancer cell lines, macrophages have been shown to have a role in metastatic seeding by performing efferocytosis of arriving tumor cells, coupled with the presence of antibodies (Kimura, et al. 2016; Thomas, et al. 2023). The role of macrophages in liver metastasis seems to be complex, as newly recruited macrophages have been shown to support metastasis and promote local immunosuppression (Peranzoni, et al. 2018; Quaranta, et al. 2018; Yu, et al. 2021). Neutrophils, on the other hand, have been shown to be recruited early upon liver metastasis, but although they have been shown to be able to control metastasis in other metastatic organs (Szlachetko, et al. 2025), their role in liver metastasis is unclear and both neutrophils and neutrophil extracellular traps have been shown to promote metastatic nidification and progression (Pearson, et al. 2025; Teijeira, et al. 2020; Yang, et al. 2020). Metastatic liver control by effector cytotoxic cells strongly relies on innate lymphoid cells like type I innate lymphoid cells (ILC1), natural killer (NK) and NK T cells (Ducimetiere, et al. 2021) and γδ T cells (Rozalen, et al. 2025). However, the role of T cells in controlling liver metastasis is controversial (Wu, et al. 2024a), given the capacity of the liver to suppress T cell responses (Hochst, et al. 2012; Limmer, et al. 2000) and the suppressive properties of macrophages and other cells in the liver (Beckinger, et al. 2023; Quaranta, et al. 2018). Interestingly, CD4^+^ T cell polarization and spatial localization determine the outcome of colorectal liver cancer and TGF-β promotes colorectal liver metastasis by inhibiting T cell responses (Tauriello, et al. 2018).

The generation of effective CTLs with the ability to recognize and kill tumor cells requires a particular form of antigen presentation termed cross-presentation. In this process, cell-associated antigen acquired from infected, or in this case, cancerous cells, rather than being presented in MHC class II to CD4^+^ T cells, can enter into a differential route that leads to loading on MHCI molecules to be directly presented to CD8^+^ T cells (Nierkens, et al. 2013). Antigen cross-presentation is mainly performed by conventional dendritic cells type I (cDC1) (Luri-Rey, et al. 2025). cDC1s are particularly important for the onset of anti-tumor CTLs (Hildner, et al. 2008) but also in anti-cancer functions beyond crosspriming of CTLs (Bottcher and Reis e Sousa, 2018) and are required for the effectiveness of many types of immunotherapies (Lopez-Janeiro, et al. 2026; Teijeira, et al. 2022). Only emerging evidence suggests that cDC1 contributes to CTL activation during metastasis (Headley, et al. 2016; Zhou, et al. 2020).

Here, we take advantage of intravital microscopy in the liver of multiple immune cell reporter mice to reveal a stepwise immune response occurring in the liver receiving cancer cells that encompasses both innate and adaptive responses. Single-cell transcriptomics, antigen cross-presentation assays and in vivo assessment of cell-cell interactions uncover active anti-metastatic surveillance by local and recruited macrophages and cDCs, which organize immune clusters to enable effective control of liver metastasis, which is therapeutically actionable by Flt3L-mediated expansion of cDCs.

## METHODS

### Cell Lines

The murine colon adenocarcinoma cell line MC38 was kindly provided by Dr. Karl E. Hellström (University of Washington, Seattle, WA) in September 1998. B16OVA cells, a murine melanoma cell line stably expressing ovalbumin (OVA), were generously provided by Dr. Lieping Chen (Yale University, New Haven, CT) in November 2001. B16F10 cells, a metastatic variant of murine melanoma, were purchased from the American Type Culture Collection (ATCC) in June 2006. The murine lung adenocarcinoma cell line CMT167 β2-microglobulin knockout and OVA-expressing cells were generated in the laboratory of Dr. Alfonso Calvo. MC38 and B16OVA-PUFFFINmCH cells were generated by Lipofectamine 3000 (Thermo Fisher) transfection with pCAGGs-mCherry (a kind gift from Dr. Monsterrat Arrasate CIMA, Pamplona) and the PUFFFINmCH (a kind gift from Sally Lowell (Lebek, et al. 2024) (University of Edinburgh, UK)) expression plasmid and selected based on cell sorting in a MoFloAstrios EQ (Beckman Coulter). All cancer cell lines, including MC38, B16OVA, B16F10, CMT167-β2m-KO-OVA, and B16OVA-β2m-KO, were maintained in Dulbecco’s Modified Eagle Medium (DMEM; Gibco) supplemented with 10% heat-inactivated fetal bovine serum (FBS; Sigma-Aldrich), 50 µmol/L β-mercaptoethanol, 100 U/mL penicillin, and 100 µg/mL streptomycin (Gibco). To maintain OVA expression in the B16OVA cell line, 400 µg/mL geneticin (Gibco) was added to the culture medium. Cells were routinely passaged by washing with PBS to remove residual serum, followed by trypsinization.

All cell lines were cultured in a humidified incubator at 37°C with 5% CO₂ for at least 7 days before in vivo inoculations. Mycoplasma contamination was routinely monitored every 8 weeks using the MycoAlert Mycoplasma Detection Kit (Lonza).

### Mice

All animal procedures were conducted in accordance with Spanish regulations for animal experimentation and were approved by the Ethics Committee for Animal Experimentation of the Regional Government of Navarra (studies 089/24, 004/26). Mice were housed at the animal facility of the Cima Universidad de Navarra (Pamplona, Spain) under specific pathogen-free conditions, with a 12-hour light–dark cycle and controlled temperature. Female C57BL/6 mice (6 weeks old) were purchased from Envigo (Barcelona, Spain). WDFY4^−/−^ (Wdfy4^em1Cya^) (Theisen, et al. 2018), *Ncr1*^GFP/+^, OT-I, Kikume (C57BL/6-^Tg(UBC*-KikGR)3-1Bhr/JI^), Foxp3EGFP, CD11cEGFP and EGFP (C57BL/6-^Tg(CAG-EGFP)1Osb/J^) mice were obtained from Jackson. XCR1-DTR-Venus mice (Xcr1^tm2(HBEGF/Venus)Ksho^) (Yamazaki, et al. 2013) were obtained from the RIKEN BioResource Research Center. XCR1iCre-mCherry (B6(129S4)-Xcr1^tm1.1(cre)Kmm^/J) mice were obtained from the Jackson Laboratory and crossed with uLIPSTIC mice (Nakandakari-Higa, et al. 2024)(C57BL/6-Gt^(ROSA)26Sortm1(CAG-Thy1,-srta)Vic/J^), obtained from Jackson Laboratory. Catchup mice (C57BL/6-Ly6g^tm2621(Cre-tdTomato)Arte^) were a kind gift from Mathias Gunzer (Hasenberg, et al. 2015), and hCD2dsRED (named *CD2^RFP^* throughout the paper) mice were a kind gift from Mark Coles from the University of York (Veiga-Fernandes, et al. 2007). Heterozygous OTIxCD2RFP, CD11cEGFPxCD2RFP and Foxp3EGFPxCD2RFP were generated by crossing these strains and F1 mice were used for experiments.

### Liver metastasis in vivo modelling

Cultured tumor cells were harvested using trypsin-EDTA before reaching confluence and resuspended in phosphate-buffered saline (PBS). A total of 5×10^5^ tumor cells were injected subcutaneously (s.c.) in 50 μL PBS using 29G syringes into the shaved right flank of ketamine– xylazine-anesthetized 6– to 12-week-old mice.

7 to 9 days later, mice were subjected to surgery. Briefly, mice were anesthetized with isoflurane, injected s.c. with 0.05 mg/kg buprenorphine, shaved and sprayed with antiseptic. A small laparotomy was performed to allow the externalization of the spleen. The spleen was gently exteriorized onto sterile, saline-moistened gauze. A single-cell suspension of 5×10^5^ CMFDA/Cell tracker Deep Red-labelled or mCherry fluorescent tumor cells was injected into the splenic parenchyma using a 29G needle. To ensure the entry of tumor cells into the portal circulation, the injection site was pressed with a sterile cotton swab for one minute to prevent leakage and ensure hemostasis. Following the injection, the splenic artery and vein were ligated at the hilum, and a total splenectomy was performed to prevent the development of a primary splenic tumor. The abdominal wall and skin were closed in two layers using V489H VICRYL 6/0 AG 11MM.

### Depletion studies and therapeutic treatments

Antibody-based depletion experiments were performed with a single-dose-based injection 24 hours prior to intrasplenic injection in all the experiments (unless otherwise indicated) for the following cell populations: 100 μg anti-CD8β (BioXCell, clone Lyt3.2) for CD8 T cells, 100 μg anti-NK1.1 (BioXCell, clone PK136) for NK cells, 100 μg anti-CD4 (BioXCell, clone GK1) for CD4 T cells, 25 μg anti-Ly6G (BioXCell, clone 1A8) + 50 μg anti-rat Kappa Immunoglobulin Light Chain (BioXCell, clone MAR18.5) for neutrophils, 400 μg anti-CSF1R (BioXCell, clone AFS98) for macrophages, 30 μg anti-CD200R3 (Biolegend, clone Ba160) for basophils. To deplete XCR1⁺ cDC1s, heterozygous XCR1-DTR-Venus mice were used. Diphtheria toxin (DT, Sigma-Aldrich) was administered intraperitoneally at a dose of 625 ng per mouse, diluted in PBS and to deplete total DCs on CD11C DTR, 200 ng of DT per mouse was administered i.p.. DT was given on days –2 and –1 before intrasplenic injection. Clodronate-based depletion was performed using 200 μl of clodronate-loaded liposomes (LIPOSOMA) and S1P-based T-cell egress from LNs was blocked with a two-dose regimen of 50 μg/mice of FTY720 (Merck) on the day of intrasplenic tumor cell injection and a day later. Pertussis toxin (Ptx treatment) was performed by intraperitoneal administration of 3 µg of Ptx (TOCRIS) in 100 µL (0.03 mg/mL) per mouse: the first dose on the day of the intrasplenic injection, followed by a second dose the next day, coinciding with the IVM session. For therapeutic experiments, a single dose of 50 μg of InVivoMAb recombinant Flt-3L-Ig (BE0342, BioXCell) or 200 μg of anti-CD47 (BioXCell, clone MIAP301) was used.

### Tissue Processing

At the specified time points, mice were euthanized using a gradually increasing concentration of CO₂. To isolate hepatic single-cell suspensions, we utilized the gentleMACS Octo Dissociator with Heaters and the Liver Perfusion Kit (Miltenyi Biotec) according to the manufacturer’s instructions. Briefly, the left lateral lobe was resected, placed into a gentleMACS Perfuser, and subjected to automated pre-digestion and enzymatic perfusion cycles at 37°C using a combination of Enzymes D, R, and A. The softened tissue was then transferred to a gentleMACS C Tube for mechanical dissociation using the LIPK_HR_1 program. The resulting homogenate was filtered through a 100 µm SmartStrainer, and the cell suspension was fractionated via differential centrifugation: a low-speed spin at 50 × g for 3 minutes to pellet parenchymal hepatocytes, followed by a 167 × g spin of the supernatant for 7 minutes to recover the non-parenchymal cell (NPC) fraction for subsequent downstream analysis.

Liver draining lymph nodes (DLNs) were mechanically disaggregated using gentle mechanical disruption and subsequently passed through a 70 µm cell strainer (Thermo Fisher Scientific) to obtain single-cell suspensions. The resulting cell suspensions were then washed with PBS and immediately stained for flow cytometry analysis.

### Flow cytometry

Surface staining for flow cytometry was performed in PBS containing viability dyes, either Zombie NIR (BioLegend, 423106) or IR808 (Invitrogen), and Fc receptor blockers. TruStain FcX anti-mouse CD16/32 block (clone 93; BioLegend, Cat# 101320, RRID:AB_1574975) was used together with the corresponding antibody mixes. Cells were then stained for 10 minutes at 4°C using fluorophore-conjugated primary antibodies against the following targets: CD11b (clone M1/70; BioLegend, Cat# 101216, RRID:AB_312799), CD11c (clone N418; BioLegend, Cat# 117334, RRID:AB_2562415), CD137 (clone 17B5; BioLegend, Cat# 106110, RRID:AB_2564297), CD172a (SIRPα) (clone P84; BioLegend, Cat# 144013, RRID:AB_2564060), CD19 (clone 6D5; BioLegend, Cat# 115530, RRID:AB_830707), CD200R3 (clone Ba13; BioLegend, Cat# 142205, RRID:AB_2144577), CD25 (clone PC61; BioLegend, Cat# 102043, RRID:AB_2561998), CD4 (clone GK1.5; BioLegend, Cat# 100451, RRID:AB_2562600 and BD Biosciences, Cat# 570257, RRID:AB_2943260), CD44 (clone IM7; BD OptiBuild, Cat# 570613, RRID:AB_2943261), CD45 (clone 30-F11; BioLegend, Cat# 103137, RRID:AB_2563312 and Cat# 103114, RRID:AB_312979), CD45.2 (clone 104; BioLegend, Cat# 109806, RRID:AB_313444), CD49b (clone DX5; BioLegend, Cat# 108922, RRID:AB_2130722), CD8α (clone 53-6.7; BioLegend, Cat# 100737, RRID:AB_2562506), CD8β (clone KT15; MBL, Cat# D271-4, RRID:AB_591031), CD8β (clone YTS156.7.7; BioLegend, Cat# 126616, RRID:AB_10642106), F4/80 (clone BM8; BioLegend, Cat# 123117, RRID:AB_893492 and Cat# 123110, RRID:AB_893490; clone T45-2342; BD OptiBuild, Cat# 750644, RRID:AB_2744383), I-A/I-E (clone M5/114.15.2; BD Biosciences, Cat# 570619, RRID:AB_2943262 and BioLegend, Cat# 107626, RRID:AB_2191071), Ly6G (clone 1A8; BioLegend, Cat# 127626, RRID:AB_2563263, Cat# 127610, RRID:AB_1877164, and Cat# 127645, RRID:AB_2565881), CD161 (NK1.1) (clone PK-136; Invitrogen, Cat# 47-5941-82, RRID:AB_1272212), PD-1 (clone 29F.1A12; BioLegend, Cat# 135220, RRID:AB_2563683), XCR1 (clone ZET; BioLegend, Cat# 148216, RRID:AB_2715797), and PE-conjugated anti-Biotin (clone REA746; Miltenyi, Cat# 130-110-951, RRID:AB_2725946). For cell sorting, liver single cell suspensions were stained with a negative lineage (CD3, NK1.1, B220, CD8β) and a positive lineage for myeloid cell populations (CD11b, F4/80, I-A/I-E). Data were acquired using a CytoFLEX LX or S flow cytometer from Beckman Coulter.

For imaging flow cytometry experiments and cell sorting, a BD FACSDiscover™ spectral flow cytometer (BD Biosciences) was used.

### Intravital microscopy of the liver

Intravital imaging of the liver was performed as previously described (Marques, et al. 2015). Mice were anesthetized with 2.5% isoflurane for induction and maintained at 1.8% during the procedure. The liver was surgically exposed via a midline and subcostal incision. To minimize motion artifacts, the rib cage was retracted using surgical thread, and the liver lobe was stabilized against a coverslip using saline-moistened tissue paper. Core body temperature was maintained at 37°C using a rectal probe-controlled heating blanket (Kemp). When indicated, mice were injected with 2,5 μg of AF647-conjugated anti-F4/80, CD45, MHCII, CD8, CD4 or anti-CD4, AF488-conjugated anti-CD4, F4/80 antibodies just before the start of surgical exposure of the liver. To define hypoxic regions, mice were i.v. injected with 20 μl of HypoxiSense™ 680 probe (Perkin Elmer) 1 hour prior to imaging.

Imaging was conducted using an LSM880 inverted microscope (Zeiss) equipped with a 25X water immersion objective (NA 0.8). Mice were secured in a custom-built stage, and z-stacks were selected based solely on the presence of tumor cells to ensure unbiased sampling. Imaging sessions lasted 3–4 hours, with time-lapse sequences acquired over 60-120 minutes at 30–60 second intervals.

Analysis of images and videos was performed using IMARIS V10 and V11 software. Neutrophils, NK cells and T cells were segmented as spots, while dendritic cells and macrophages were segmented using the surface tools. In CD2RFP segmentation, only cells with high RFP expression levels were included. Segmented T, NK cells and neutrophils were semi-automatically tracked using an autoregressive motion algorithm; tracks corresponding to improperly segmented cells and tracks present on the borders of the fields were removed for the analysis. Tracks were manually corrected by the user for mistakes in cell movement assessments. Dendritic cells were manually tracked by the users. F4/80 cells were not normally tracked. For behavioral analysis, tracks were selected based on total track length (>100 μm patrolling, <100 μm stopping). To define the interaction between immune cells, spots with a distance < 3 μm to a macrophage/DC surface were defined as contacting and spots with a distance > 3 μm were defined as not contacting. In some experiments, tracking was performed within different regions within the same images.

### Adoptive T-cell transfer assays

Total splenocytes were isolated from spleens and lymph nodes of OTI x CD2RFP by mechanical disruption and activated with 1ng/ml SIINFEKL peptide with 25 U/ml rhIL-2 (Proleukine) for 48h. Then, the cells were centrifuged, and the media with IL-2 were refreshed for another 48 hours.

Expanded, activated CD8⁺ T cells were labeled with 5 μM CellTrace Violet (CTV, BioLegend) at 37 °C for 15 minutes and washed twice with PBS prior to transfer when proliferation was to be analyzed by flow cytometry. 4×10^6^ activated OTI T cells were injected intravenously 24-48h before tumor intrasplenic injection as indicated in the figures. The proliferation index (PI) of CD8^+^ T lymphocytes labeled with CTV was calculated using the formula PI = [Σ (fₙ × 2ⁿ)] / [Σ fₙ], where *f*n represents the fraction of cells that have undergone n divisions, and 2^n^ reflects the theoretical cell expansion resulting from those divisions.

### Label of cDC1-T cell Interactions *In Vivo* using the Xcr1-Cre-uLIPSTIC System

To assess in vivo cell–to–cell interactions involving cDC1, the XCR1uLIPSTIC mouse model, crossed in our laboratory, was used. Mice were subcutaneously injected in the right flank with 5 × 10⁵ MC38 tumor cells. On day 7 post-tumor implantation, mice were injected i.s. with 0.5 × 10⁵ tumor cells. Twenty-four hours later (day 8), mice received intraperitoneal injections of the biotinylated transpeptidase substrate (biotin–aminohexanoic acid–LPETG) every 20 minutes for 2 hours (100 µL of a 20 mM solution in PBS per injection). 20 minutes after the final injection, mice were sacrificed, and livers and liver DLNs were collected and processed for flow cytometry analysis to detect biotin transfer to CD90.1 as a readout of cDC1-mediated interactions with T cells.

### Tumor to liver migration assessment in Kikume green photoconverted mice

To study T cell migration from the primary tumor to the liver, we used Kikume green mice to s.c. implant MC38 tumors in the right flank as previously explained. At day 7 post s.c. tumor injection, mice were subjected to i.s. tumor cell or PBS injection followed by splenectomy. At day 8, s.c., tumors were photoconverted. To do so, mice were anesthetized with ketamine and xylazine, the tumor area was shaved with Veet cream, and animals were covered with aluminium paper except for the s.c. tumor present in the back and subjected to 10 minutes of irradiation with a 405nm LED light at 150 mW of power. Such a treatment typically reached 40% –60% of photoconversion of stromal cells from KikG to KikR as assessed by flow cytometry. 24 hours later, mice were sacrificed and livers analyzed by flow cytometry. To set single staining controls for the proper assessment of photoconversion, a fraction of a spleen single cell suspension from a non-photoconverted mouse was subjected to 1-minute UV irradiation to reach 100% photoconversion and another fraction of the same single cell suspension was kept in the dark.

### Liver tissue clarification for imaging

Pieces of liver of around 5 mm^3^ were collected and put into 2 mL microcentrifuge tubes containing 500 μL of Antigen Fix (Diapath) and incubated for 24 hours at 4°C. Following fixation, tissue was transferred to 300 μL of FDC clarification buffer (phosphate-buffered saline [PBS] supplemented with 1% BSA, 2% Mouse serum, and 2% Triton X-100). For long-term storage, samples were maintained in this buffer at 4°C. For whole-mount labeling, livers were incubated with AF647-conjugated anti-F4/80 antibodies (1:200) in FDC clarification buffer for 3–4 days at RT under continuous agitation. Following staining, tissues underwent a 24-hour wash in FDC clarification buffer to remove unbound antibodies. To achieve optical transparency, liver pieces were transferred to 2 mL microcentrifuge tubes containing 300 μL of RapiClear 1.52 (Refractive Index [RI] = 1.52; SunJin Lab). Samples were incubated in RapiClear until cleared. Cleared tissues were stored at 4°C or maintained at RT for 24 hours prior to imaging to ensure complete RI equilibration. Cleared liver pieces were mounted in RapiClear solution within 35 mm polymer-bottom dishes (IBIDI). 3D fluorescence imaging was performed using an LSM880 confocal microscope (Zeiss) equipped with a 25X water immersion objective (NA 0.8).

### Raw data processing for scRNAseq analysis

Raw sequencing reads from each BD Rhapsody™ library (Positive and Negative conditions) were processed independently using the BD Rhapsody Sequence Analysis Pipeline v2.3, executed via CWL workflow. Reads were aligned to the mouse GRCm39 reference genome using the prebuilt BD WTA reference RhapRef_Mouse_WTA_2025-10, which incorporates the GENCODE vM38 primary assembly annotation (gencode.vM38.primary_assembly.annotation-processed.gtf).This pipeline performs read trimming, STAR-based alignment, UMI deduplication, and cell-barcode calling, producing per-cell gene expression matrices in MEX format for both the filtered (cell-containing) and unfiltered (all droplets) outputs. Libraries were sequenced on a NextSeq 2000 platform.

### Cytokine and chemokine profiling in liver homogenates using ProCartaPlex Multiplexing

Cytokine levels in the liver were quantified using a customized ProCartaPlex Multiplex Immunoassay (Thermo Fisher Scientific). Briefly, liver samples were thawed at room temperature and subsequently resuspended in 150 µL of RIPA buffer (Tris 65 mM, KCl 150 mM, Np40/IGEPAL 1%, SDS 0.1%, pH 7.4) supplemented with a cocktail of protease (cOmplete Mini, Roche) and phosphatase (PhosSTOP, Roche) inhibitors. The tissue was then mechanically homogenized using a pellet mixer. To isolate the protein fraction, the resulting homogenate was centrifuged at 75,000 rpm for 20 minutes at 4°C. The supernatant containing the liver proteins was collected. Cytokine concentrations were measured according to the manufacturer’s instructions using a Luminex MAGPIX. Data were normalized to the total protein concentration of each sample, as determined by a BCA assay (Thermo Fisher Scientific).

### Ambient RNA correction and doublet detection

All downstream analyses were performed in R (v4.3) using Seurat (v5) (Hao, et al. 2024). Ambient RNA contamination was estimated and corrected independently per sample using SoupX (v1.6.2) (Young and Behjati 2020). The unfiltered and filtered MEX matrices were loaded with Read10X. A temporary Seurat object was constructed from the filtered matrix, processed through standard normalization, dimensionality reduction, and Louvain clustering (resolution = 0.5) solely to provide cluster labels to the SoupX contamination model. The contamination fraction was estimated automatically per cluster with autoEstCont and per-gene, per-cell corrected integer count matrices generated with adjustCounts(roundToInt = TRUE).

From each SoupX-corrected matrix, a Seurat object was created with CreateSeuratObject (min.cells = 3, min.features = 200). Doublet detection was performed with scDblFinder (v1.14; clusters = TRUE, set.seed = 123) (Germain, et al. 2021). Cells classified as doublets were removed prior to all downstream analyses.

### Quality control and cell filtering

Cells were filtered based on three quality metrics: number of detected genes (nFeature_RNA), total UMI count (nCount_RNA), and the fraction of reads mapping to mitochondrial genes (percent.mt). Cells were retained if they simultaneously satisfied: nFeature_RNA > 200 and < 5,000; nCount_RNA > 500 and < 40,000; percent.mt < 20%. These thresholds were deliberately permissive to preserve granulocyte populations — neutrophils and basophils — which physiologically display lower transcriptomic complexity and elevated mitochondrial transcript fractions compared to monocyte–macrophage populations.

### Data integration, dimensionality reduction, and clustering

Seurat objects from both conditions were merged into a single object with merge, preserving condition metadata. Data were log-normalized (NormalizeData) and the top 2,000 highly variable genes identified with FindVariableFeatures. Data were scaled regressing out library size and mitochondrial fraction (ScaleData, vars.to.regress = c(“nCount_RNA”, “percent.mt”)). PCA was run with RunPCA (npcs = 50) and the first 20 principal components retained based on elbow plot inspection. Neighbor graph construction (FindNeighbors, dims = 1:20, algorithm = 4 [Leiden]), community detection at resolutions 0.5, 0.8, and 1.0 (FindClusters), and UMAP embedding (RunUMAP, dims = 1:20) were performed on the PCA embeddings directly.

### Cell type annotation

Automated cell type annotation was performed with SingleR (Aran, et al. 2019) using the murine steady-state liver myeloid atlas from Guilliams et al. as a reference (Guilliams, et al. 2022). This atlas comprises 71,261 myeloid cells annotated into 11 cell type categories: Kupffer cells, Ly6C^hi^ classical monocytes, non-classical monocytes, monocyte-derived macrophages (MoMFs), CD74^+^ MHC-II^hi^ macrophages, lipid-associated macrophages (LAMs), inflammatory macrophages, cDC1, cDC2, plasmacytoid dendritic cells, and neutrophils. The reference atlas was log-normalized prior to use. SingleR was executed in per-cluster mode, assigning labels based on Spearman correlation between cluster-average expression profiles and reference cell type signatures.

A second annotation pass was performed using the complete Guilliams liver atlas (all hepatic cell types) to flag non-myeloid contaminating cells captured during sorting. Cells annotated as hepatocytes, endothelial cells, liver sinusoidal endothelial cells (LSECs), stellate cells, cholangiocytes, fibroblasts, mesothelial cells, T cells, B cells, NK cells, or ILC1s were excluded from all downstream analyses.

The broad “Monocytes & Monocyte-derived cells” atlas category was resolved at higher resolution. Cells belonging to this category were extracted and re-processed independently: log-normalization, FindVariableFeatures (nfeatures = 2,000), ScaleData, RunPCA (npcs = 30), RunUMAP (dims = 1:20), FindNeighbors (dims = 1:20), and FindClusters (resolution = 0.5), yielding 13 clusters. Cluster identity was assigned by manual annotation based on average expression of canonical marker genes: Ly6c2, Ccr2, S100a8, S100a9 (Ly6C^hi^ classical monocytes); Cx3cr1, Csf1r (intermediate monocytes); Cd74, H2-Aa, H2-Ab1 (CD74^+^ MHC-II^hi^ macrophages); Cxcl9, Cxcl10, Ccr5 (inflammatory macrophages); Apoe, Trem2, Spp1 (LAMs/foamy macrophages); and Mki67, Top2a (proliferating monocytes). Five transcriptionally convergent clusters (mean Cd74 > 3.0 and H2-Aa/H2-Ab1 > 2.0 in log-normalized expression) were merged into a single CD74+ MHC-II^high^ Macrophage population, yielding nine final monocyte/macrophage subpopulations.

### Annotation correction by spatial re-assignment

During quality review of the joint UMAP, subsets of cells annotated as Kupffer cells and neutrophils were found to be spatially dispersed within UMAP regions dominated by monocyte– macrophage clusters, inconsistent with their expected transcriptomic neighborhood. This pattern is characteristic of low-confidence SingleR label transfer, where cells with ambiguous expression profiles receive the highest — yet still low — correlation score to an incorrect reference category. Spatially discordant cells were identified using UMAP coordinate thresholds (Kupffer cells: UMAP_2 < −7; neutrophils: UMAP_2 < −5) and re-assigned by majority vote among their 30 nearest neighbors (k = 30, FNN v1.1.4.1), restricted to the nine monocyte/macrophage reference populations. Cells falling within the principal UMAP islands — forming compact, spatially coherent clusters with unambiguous canonical marker expression (Timd4, Vsig4, Clec4f for Kupffer cells; Csf3r, Cxcr2, S100a8, S100a9 for neutrophils) — were retained with their original annotation irrespective of SingleR confidence score. The final annotated object comprised 39,677 cells across 16 myeloid populations.

### Neutrophil subclustering using the NeuMap atlas

Neutrophils were extracted from the final annotated object (n = 4,928; 4,628 Negative, 300 Positive) and re-processed independently: log-normalization, FindVariableFeatures (nfeatures = 2,000), ScaleData, RunPCA (npcs = 30), RunUMAP (dims = 1:20), FindNeighbors (dims = 1:20), and FindClusters (resolution = 0.5). Cluster identity was assigned based on expression of NeuMap hub markers (Cerezo-Wallis et al., 2026): Immature/PreNeu (Mmp8, Ngp, Camp, Ltf), IFN-response (Ifit1, Isg15, Irf7, Rsad2, Stat1, Gbp2, Mx1), IS-I/Immunosuppressive (Nos2, Hilpda, Il1a, Il1rn, Ero1a, Cd14, Ptgs2, Trem1), and Antigen Presentation (H2-Ab1, Cd74, Ctss). As IS-I and IS-II states could not be reliably distinguished in this dataset, these populations were merged into a single IS-I/Immunosuppressive cluster, resulting in a final annotation of four subpopulations. Differential abundance of Positive and Negative cells across subpopulations was assessed by the chi-square test.

### Differential gene expression analysis and visualization

Differential gene expression (DEG) between Positive and Negative conditions was performed independently for Kupffer cells, CD74⁺ MHC-II^^high^^ Macrophages, and Intermediate Monocytes using FindMarkers (Wilcoxon rank-sum test, min.pct = 0.1, logfc.threshold = 0.25, Bonferroni-corrected p_adj < 0.05). DEG was not performed for cDCs (49 Positive vs. 3,999 Negative cells) or neutrophils (∼314 Positive cells — insufficient for reliable comparison; NeuMap proportions and UMAP visualizations were used instead). Results were visualized as volcano plots with genes colored by functional category, significance threshold at p_adj = 0.05, and fold-change lines at |log2FC| = 0.5; only genes exceeding this threshold were labeled.

### Curated gene expression heatmaps

Heatmaps displaying the log2 fold-change (Positive vs. Negative) of statistically significant differentially expressed genes (Bonferroni-corrected p_adj < 0.05) were generated with pheatmap (v1.0.13). Gene lists were organized into functional modules and validated by the study immunologist (A.T.S.). The following categories were defined for the Kupffer cell heatmap: Antigen Presentation (H2-D1, H2-DMb1, B2m, Cd74, Anpep, Ms4a7), Actin Dynamics (Actb, Arpc2, Arpc5, Capg, Cfl1, Cttn, Myo1e, Cdc42, Rac2, Rhoc), Antigen Uptake (Marco, Itgax), Immune Activation (Cxcl9, Cxcl16, Slamf7, Slamf8, Slamf9), Lysosomal Trafficking (Ctsz, Cd63, Atp6v0c, Atp6ap2, Mpeg1, Kif3b, Kif3c, Rab10), Survival (Bcl2a1a, Bcl2a1b, Bcl2a1d), Metabolism (Gapdh, Pgk1, Eno1, Atp5f1b, Atp5mk, Cyc1, Cox17, Lpl), Tissue Remodeling (Mmp12, Mmp13, Dnase1l3), Loss of Kupffer Identity (Clec4f, Vsig4, F13a1, Apoe), Loss of Inflammation (Ccr2, Ccl6, S100a8, S100a6, Chil3, Il1r2), and Loss of Adhesion (Alcam, Itga4, Sema4d, Vcan). Equivalent functional categories were defined for the CD74⁺ MHC-II^high^ Macrophage and Intermediate Monocyte heatmaps, with gene lists adjusted per population. Color scales were implemented asymmetrically, anchoring zero to white with independent break sequences and color ramps for positive and negative log2FC ranges, to prevent artificial color centering in datasets with asymmetric expression ranges.

### cDC1 subclustering analysis

Given the near-complete depletion of cDCs in the Positive condition (cDC1s: 1,648 Negative / 32 Positive; cDC2s: 2,319 / 17; Mig. cDCs: 81 / 0), DEG between conditions was not feasible. cDC1s were instead extracted and re-processed independently (log-normalization, FindVariableFeatures nfeatures = 2,000, ScaleData, RunPCA npcs = 15, RunUMAP dims = 1:15, FindNeighbors dims = 1:15, FindClusters resolution = 0.3), yielding two clusters. Enrichment of Positive cells per cluster was assessed by Fisher’s exact test. Differential gene expression between the Positive-enriched cluster and all remaining cells was performed with FindMarkers() (Wilcoxon, min.pct = 0.25, logfc.threshold = 0.25, p_adj < 0.05).

### Immunohistochemistry (IHC) staining and analysis

XCR1 (clone D2F8T, dilution 1:50, Cell Signaling, 44665) IHC staining and CD8 (clone C8/144B, ready-to-use, Agilent, GA62361-2) IHC staining were performed on a Roche Ventana BenchMark ULTRA Slide Stainer (Ventana Medical Systems, Inc. USA) platform with the OptiView DAB IHC detection kit. Healthy tonsil tissue sections served as positive tissue controls. Whole stained slides were scanned on the VENTANA DP 200 slide scanner. Densities of XCR1+ and CD8+ cells were quantified using the open-source digital pathology software QuPath version 0.4.4 and expressed as the number of cells per mm2.

### Quantification and statistical analyses

All data were analyzed using GraphPad Prism version 8.0. Flow cytometry data were processed with FlowJo v11 or CytExpert software.

Results are presented as dots for individual mice/cells plus median, unless otherwise indicated. Comparisons between two groups were performed using the non-parametric Mann-Whitney test unless otherwise indicated. Exact p-values are shown for any analysis.

## RESULTS

### Liver sinusoids are surveilled by discrete leukocyte subsets

The liver contains a vast vascular network composed of sinusoidal endothelial cells populated by intravascular resident macrophages (Kupffer cells) (Guilliams, et al. 2022). We hypothesized vascular immunosurveillance by non-resident myeloid and lymphoid cells might involve vascular patrolling mechanisms (Westhorpe, et al. 2018) that could be further enhanced upon the arrival of metastatic tumor cells. We therefore performed intravital imaging of the liver tissue in mice bearing a highly immunogenic subcutaneous tumor (MC38) to understand immune surveillance patterns of different immune cells in the liver pre-metastatic niche. Immune cell reporter mice (CD2RFP for T cells, NCR1 GFP for NK cells, CD11c GFP for DCs) were implanted subcutaneously with MC38 tumor cells, and their livers were imaged by multiparametric intravital microscopy.

We found that while many T cells flowed rapidly within vessels, others were retained within the liver, exhibiting intravascular patrolling behaviors, with extended crawling, change in direction, and longer residence times in the sinusoids as compared with other stationary T cells) We categorized T-cell behaviors into three different classes, depending on the residence time and distance travelled within the sinusoids. Cells that stopped for at least two frames (60 sec) but had paths shorter than 100 μm represented the majority of observed cells and were defined as “stopping” cells. Cells that stopped but showed tracks larger than 100 μm, frequently crawling within liver sinusoids, were categorized as patrolling cells (Figure 1A). Patrolling cells represented a smaller fraction of T cells, frequently ranging from 10 to 20% of total observed T cells (Figure 1B and 1C). Individual-track analysis of T-cell behavior showed that patrolling T cells travelled much longer distances than stopping T cells, had higher speeds within the sinusoids, remained for longer periods within the liver, and exhibited meandering behavior with markedly less directional migration, characterized by a reduction in their chemotactic index (Figure 1C).

**Figure 1.**
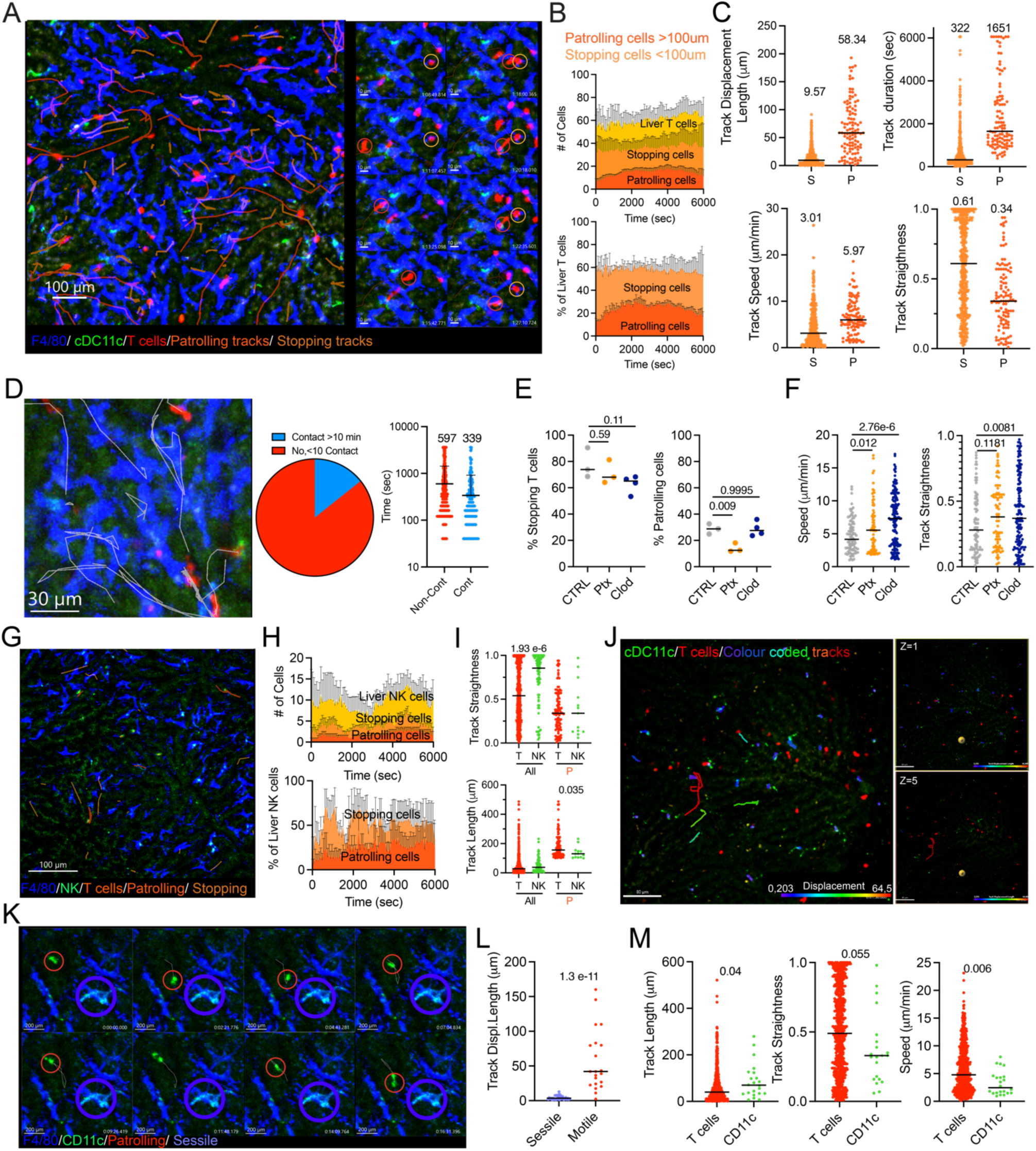
Immune surveillance of the liver sinusoids by effector immune cells and antigen-presenting cells. A) Representative images showing individual T-cell tracks of patrolling (>100 μm length), stopping (<100 μm length) T cells on a premetastatic liver. A sequence shows in colored circles representative T cells during stopping or patrolling behaviors. (B) Quantification of T cell motility patterns within the liver, expressed as both absolute cell counts and as a percentage of the total T-cell population per imaging frame. (C) Quantitative assessment of motility features of tracked T cells. Mean values are shown for each parameter and defined behavior (S, stopping; P, Patrolling). (D) Sequence depicting a patrolling T cell (white circled) interacting with local macrophages and the spatial correlation of individual T cell tracks with local macrophages. Quantification of T cells performing or non-performing stable contacts (>10 min) with local macrophages in the liver. Mean maximum residency time of single T cells within the liver, contacting or not contacting with macrophages. Mean contacting and non-contacting time values are depicted in the graph. (E) Proportion of recorded T cells displaying stopping or patrolling behaviors upon clodronate-based macrophage depletion or Ptx treatment. (F) Comparative assessment of motility features of T cells upon clodronate-based macrophage depletion or Ptx treatment. (G) Representative image of liver NK cells and their tracks characterized in Ncr1 GFP reporter mice. (H) Quantification of NK cell motility patterns within the liver, expressed as both absolute cell counts and as a percentage of the total NK cell population per imaging frame. (I) Comparative assessment of motility features of overall and patrolling T and NK cells. (J) Representative image of liver CD11c cells and their tracks color-coded by overall displacement and single Z sections of liver (Superficial, Glisson’s Capsule; and deep, parenchymal) on CD11c EGFP-CD2 RFP double reporter mice. (K) Sequence depicting the two types of CD11c cells’ motility behavior, sessile (purple circled) and motile (red circled). (L) Overall displacement of tracked sessile and motile CD11c cells within the livers. (M) Comparative assessment of motility features of overall and patrolling T and DCs. Data were analyzed from at least three videos from two different mice per condition. Each dot represents a single cell, except in E, where it represents an analyzed video. Medians are represented as a horizontal bar for each condition. Exact p-values are shown for statistical comparison, U Mann-Whitney tests and ANOVA followed by Dunnett’s T test.

Concomitant imaging of liver T cells and macrophages revealed that patrolling T cells were frequently engaging with resident macrophages within the liver sinusoids (Figure 1D). Upon interaction, T cells frequently changed their direction and maintained stretched contact with liver macrophages (Figure 1D). Quantitative assessment of this behavior showed that approximately 10% of all traceable T cells spent at least 10 minutes in contact with liver macrophages (Figure 1D). When comparing non-contacting and contacting times of individually tracked T cells, we observed that the average amount of time that a T cell could be interacting with a macrophage was around 5 minutes, a time that would allow for effective antigen scanning in the liver and comparable to T cell-DC scanning behaviors reported in mouse lymph nodes (LNs) (Miller, et al. 2004).

To understand to what extent macrophage scanning was behind the specific patrolling behavior of T cells, we performed clodronate-based depletion of liver macrophages and intravital imaging. The proportion of stopping and patrolling T cells was unaltered in macrophage-depleted mice (Figure 1E). However, patrolling T cells showed a different behavior when macrophages were absent, which was characterized by longer distances travelled, higher speeds, and straighter paths, suggesting a macrophage involvement in liver T cell retention to allow efficient scanning for antigen over the liver resident macrophages (Figure 1F). To elucidate some of the mechanisms defining this patrolling behavior within the liver and understand if such patrolling behaviors were actively regulated, we inhibited Gi signaling using Pertussis Toxin (Ptx) to inactivate most chemotactic-related receptors on T cells. Ptx treatment did not impact the proportion of T cells performing short stops, but effectively reduced the proportion of patrolling cells, suggesting that crawling within sinusoids was a chemokine-regulated process (Figure 1E). Individual T cell track analysis evidenced straighter and faster movement of T cells, consistent with less active T cell retention within the liver (Figure 1F).

To define if the observed T cell patrolling behavior was exclusively characteristic of CD8^+^ or CD4^+^ T cells, we injected CD2RFP mice with fluorescent anti-CD8β and anti-CD4 antibodies just before imaging. Our motility analysis showed that CD4^+^ T cells were more frequent within the liver, but the proportional behavior in terms of patrolling and stopping-cell categories was similar for CD4^+^ and CD8^+^ T cells. Overall analysis of all traceable CD4^+^ and CD8^+^ T cells did not show any major differences in motility behaviors, but selective analysis on patrolling (>100 μm of displacement) T cells showed that CD4^+^ T cells tend to have a more efficient patrolling behavior, with larger distances, higher speeds and straighter paths than their CD8 counterparts (Figure S1A and 1B).

Due to the well-known tolerogenic state of the liver (Beyzaei, et al. 2022), we decided to determine whether the CD4^+^ population exhibiting strong patrolling behavior was enriched in T regulatory cells. Imaging Foxp3xCD2RFP heterozygous mice, we observed a few double-positive Foxp3^+^ CD2RFP cells being retained within the liver, in a similar proportion to other non-Foxp3^+^ T cells within the same videos (Figure S1C). These observations indicate that only a small fraction of the observed CD4^+^ T cells are T regulatory cells (Tregs). Again, the general behavior of Foxp3 positive and Foxp3 negative red fluorescent T cells was unchanged, but the few patrolling Foxp3 GFP cells caught patrolling showed a similar behavior to previously characterized CD4^+^ T cells, with higher patrolling speed and longer stays within the liver (Figure S1C).

To understand if such liver immunosurveillance, visualized as liver patrolling, was extensive to other immune cell types known to be effector cells in anti-metastatic responses, we performed similar experiments in other fluorescent reporter mouse strains. NK cells are well known to be important in metastasis control and in liver immunity (Lopez-Soto, et al. 2017). Liver live imaging of NK cells and NK T cells in an Ncr1 GFP reporter mouse in this setup identified fewer NK cells being retained in the liver at baseline, with lower proportions of NK cells showing a patrolling behavior, as compared to T cells (Figure 1G and 1H). Single-cell track analysis supported this lack of retention, characterized by much straighter cell paths in all traceable NK cells and only minor changes in patrolling NK cells (track length> 100 μm), which displayed shorter paths, unexpectedly indicating that at least in this reporter mouse, NK cells showed a more limited patrolling behavior within the liver than T cells (Figure 1I). To characterize DC behavior in the liver, we imaged CD11c reporter mice (CD11c DTR EGFP and CD2RFPxCD11cEGFP mice). Live microscopy videos clearly show the presence of a majority of non-motile DCs located either in the liver capsule or around periportal areas (Sierro, et al. 2017; Watanabe, et al. 2007). However, we could also identify some CD11c cells that were actively patrolling the liver sinusoids (Figures 1J-L). Liver patrolling CD11c cells exhibited a different behavior from that of T cells when compared in the same videos obtained in the double reporter CD11cEGFPxCD2RFP model, characterized by active crawling over the endothelium with lower speeds, longer tracks, and less directional movements, denoting a stronger patrolling behavior in these local professional APCs within the liver sinusoids as compared to T cells. Overall, we can conclude that the healthy premetastatic liver is actively surveilled and patrolled by dendritic cells, NK cells, and T cells, but that each population shows specific dynamics that may dictate their behaviors upon the appearance of pathogens or metastatic tumor cells.

### The early immune response against tumor cells arriving in the liver is characterized by neutrophil interactions and a strong macrophage phagocytic response

To define how immune responses start in the liver after the arrival of pioneering metastatic cells, we performed intravital imaging shortly (five minutes) or soon (2-4h) after injection of fluorescent MC38 tumor cells in the spleen (to allow tumor cell arrival via the portal vein) of mice bearing old subcutaneous tumors for 7 days. Live imaging focusing on tumor cell behavior (Figure 2A) revealed that, similarly to what has been described in metastasis to the lung (Headley, et al. 2016), most of the cells start to break and release large vesicles or cell pieces to the circulation upon arrival at the liver capillaries. Other tumor cells also started to extend their cytoplasm and seemed to colonize the liver vascular areas. Imaging of these steps in a mouse reporter with ubiquitous EGFP expression allowed us to identify two key events: first, that some circulating immune cells rapidly interacted with the arriving tumor cells, and second, that flow stasis could be observed in vessels populated by embolized tumor cells, with slow-flowing platelets only in close vicinity to tumor cells (Figure 2B). The speed and rapid appearance of these EGFP cells, constantly interacting with recently arrived tumor cells, suggested that these cells could be neutrophils. Imaging of the same phenomenon in the Ly6G^Tomato^ mice (Hasenberg, et al. 2015) confirmed prominent and rapid neutrophil infiltration in livers with metastatic cells. Neutrophils were characterized by rapid directional migration toward tumor cells, which was followed by “repulsion” from the tumor cell area (Figure 2C). Quantitatively, neutrophils showed enhanced retention in the metastatic liver (Figure 2D and F), with higher speeds and longer migration tracks (Figures 2E and G), suggesting cell activation. Imaging of other reporter strains showed that neither NK cells nor CD11c cells had a clear directional migration towards metastatic tumor cells (Figure S2A-F). T cells could also be found interacting with tumor cells, but lacked directional migration towards tumor cells (Figure 2H). Single cell tracking of T cells at this time point in metastasis showed no evident recruitment to the liver over time and a small drop in all motility parameters (Figure 2H-J), a phenomenon also observed in NK cells, suggesting that effector arrest in the early hours of metastatic establishment is more a passive mechanism than an active one, and that this retention is rather caused by blood flow perturbations within the liver rather than active production of chemokines by arriving tumor cells or other liver cell sources. Neutrophil recruitment, however, showed the features of an active process, with directional migration and very short interactions with metastatic tumor cells (Figure 2K), suggestive of active killing or trogocytosis mechanisms as previously described (Belz, et al. 2026; Matlung, et al. 2018).

**Figure 2.**
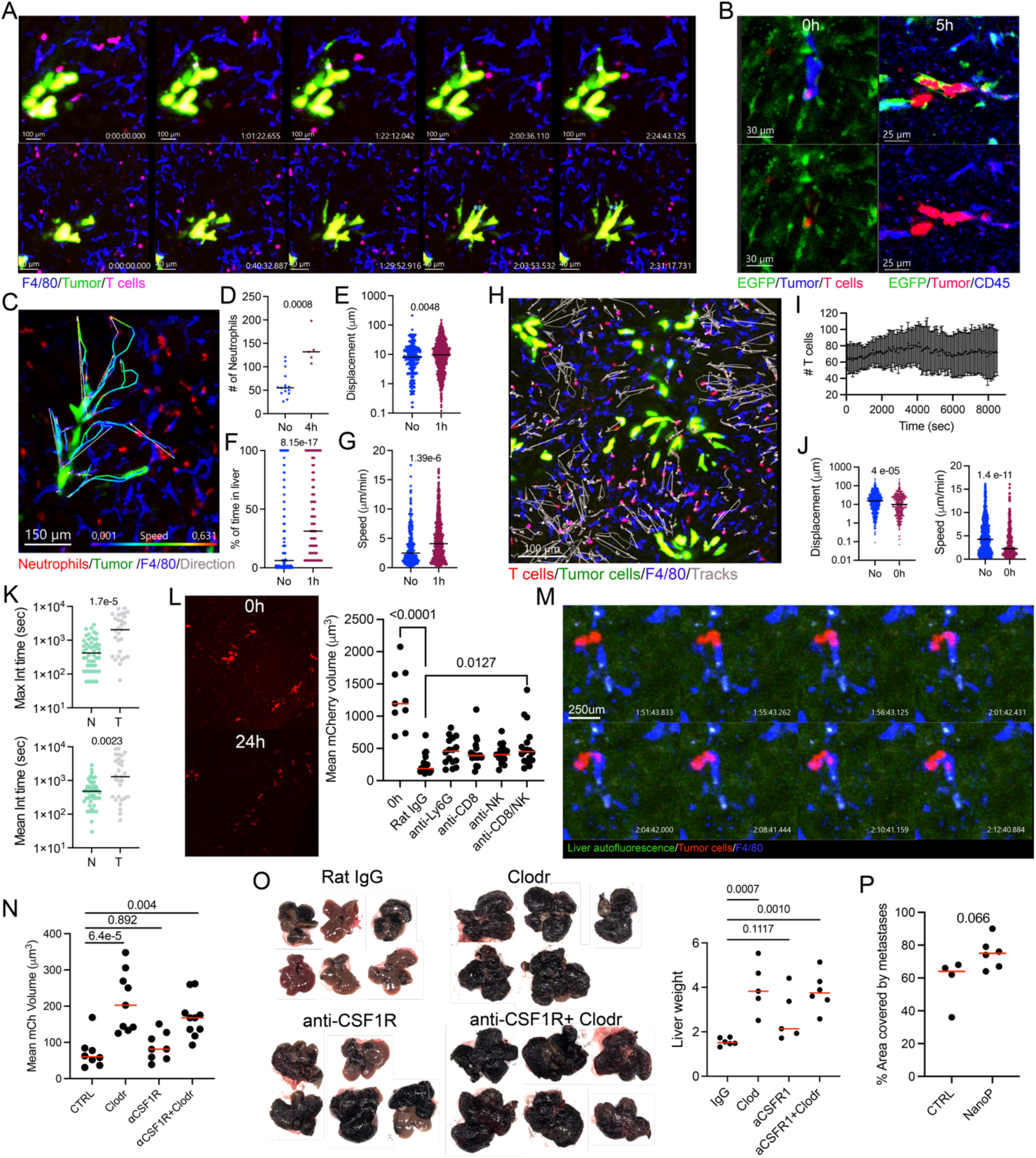
Incoming metastatic cells elicit neutrophil recruitment and macrophage phagocytosis. (A) Intravital imaging sequence showing MC38 tumor cell behavior right upon arrival to the liver. (B) Representative images of endogenous mouse cells (GFP, green) interacting with MC38 tumor cells upon arrival in the liver. (C) Representative image of neutrophil-directed migration towards tumor cells 4 hours after tumor cell arrival in Catchup mice. Selected individual tracks of neutrophils are color-coded based on cell speed and grey arrows show the direction of the overall displacement of each tracked cell. (D) Quantification of fluorescent neutrophils at the beginning of intravital microscopy videos in non-metastatic liver and at the 4h time point after intrasplenic injection. Overall displacement (E), percentage of time spent by individual neutrophils in the liver (F) and mean speed of neutrophils at the indicated conditions. (H) Representative image of T cell behavior upon metastatic cell arrival. Individual tracks of T cells are shown in white and grey arrows show the direction of the overall displacement of each tracked cell. (I) Absolute count of tracked T cells within the liver over the intravital imaging session. (J) Comparative assessment of motility features of T cells in non-metastatic liver or upon arrival of tumor cells. (K) Comparative assessment of maximum and mean interaction times of neutrophil and T cells with tumor cells. (L) Representative images of mCherry MC38 tumor cells visualized by whole liver confocal imaging at the indicated time points after tumor cell arrival (left) and quantification of the mean particle size of mCherry positive signals at the indicated time points (right) in mice pretreated or not with depleting antibodies for the indicated populations. Each dot represents a single mouse, with at least three images per mouse analyzed. (M) Intravital imaging sequence showing a macrophage performing efferocytosis of MC38 tumor cell material. (N) Quantification of the mean particle size of mCherry-positive signals in mice pretreated with the indicated compounds for macrophage depletion at the 24h post-MC38 mCh intrasplenic injection timepoint. Each dot represents a single mouse, with at least three images per mouse analyzed. (O) Representative images of the livers of mice treated with the indicated compounds prior to intrasplenic injection of B16OVA cells at day 13 post-intrasplenic inoculation, and corresponding liver weight. Each dot represents a single mouse. (P) Quantification of B16OVA liver metastatic area at day 13 post-intrasplenic inoculation in mice treated with nano-primer particles 6 hours prior to intrasplenic injection of B16OVA cells. Each dot represents a single mouse. Data were analyzed from at least three videos from two different mice per condition. Each dot represents a single cell, except in D, where it represents an analyzed video and in L, N, O, and P, in which each dot represents a single mouse (for microscopy images, three FOV per mouse analyzed). Medians are represented as a horizontal bar for each condition. Exact p-values are shown for statistical comparison, U Mann-Whitney tests (D, E, F, G, J and K), ANOVA followed by Dunnett’s T test. (L, N, O and P).

Given the numerous reports defining the role of NK cells, T cells and neutrophils in controlling metastatic spread and our observations of direct interactions between these populations, we sought to understand the relevance of these three effector populations in actually controlling the early metastatic steps. Imaging of the liver 24h after intrasplenic injection of mCherry-expressing tumor cells revealed a generalized loss of tumor cell integrity, with few cells retaining their large/intact shape at that particular time point. Red fluorescence was mainly observed forming fragments smaller than a cell in many areas of the liver at this 24h post-metastasis time point (Figure 2L). We hypothesized that leukocytes may be mediating the destruction of tumor cells through cytotoxicity, phagocytosis, or trogocytosis.

To define the contribution of different immune effector cell populations in the survival of pioneering tumor cells, we imaged the liver of mice pretreated with depletion antibodies against neutrophils, CD8^+^ T cells and NK cells 24 hours before intrasplenic tumor injection. We then quantified the size of the mCherry-positive particles to assess tumor cell integrity within the metastatic liver 24 hours later. Livers from mice treated with isotype control (Rat IgG) clearly showed a reduced mean tumor cell size at this time point when compared with those just arriving to the liver. All the depletions tested correlated with larger proportions of intact tumor cells that, in most cases, were not statistically significant and were far from the observed sizes at time 0 in the liver, indicating that these effector immune cell populations may be able to clear a portion of metastatic cells over the early anti-metastatic response, but that survival within the liver of metastatic cells was largely effector cell independent (Figure 2L).

Next, we focused on the behavior of local macrophages upon tumor cell arrival. Such cells were again visualized by staining them with an anti-F4/80 fluorescent antibody concomitantly with visualization of CD11c or T cells in the different corresponding reporter mice. Imaging macrophages showed frequent tumor material uptake, either by nibbling directly arriving tumor cells or by efferocytosis of large vesicles extruded from the tumor cells in the vascular bed (Figure 2M). To understand the importance of tumor cell engulfment by macrophages, either liver resident or newly recruited, in limiting liver metastasis, we first tried to define depleting strategies intended to differentially target both local and newly recruited macrophages. We hypothesized that clodronate treatment would mainly eliminate liver resident macrophages, while minimally impacting any other antigen-presenting populations or newly arrived macrophages. In contrast, CSF1R blockade will only partially impact local liver macrophages, but should strongly impact newly recruited macrophages, and double clodronate and anti-CSF1R treatment should affect both resident and newly recruited cells. Flow cytometry and imaging of clarified liver tissue of CD11c GFP mice treated with these depleting agents 24 hours before intrasplenic tumor injections revealed that CSF1R blockade only partially impaired the number of local macrophages, leaving some subpopulations like F4/80^Med^ MHCII^Med^ macrophages barely unchanged (Figure S2G). Clodronate almost completely removed all liver macrophages with a minor impact on cDCs, particularly those found in the parenchyma of the liver, while the double depletion generated a more pronounced elimination of all phagocytic subsets (Figure S2G and S2H). In this setting, both clodronate and combined anti-CSF1R plus clodronate treatment were associated with significantly higher numbers of intact tumor cells in the liver, whereas the effects of single-agent CSF1R blockade were only partial and not significant (Figures 2N and S2I). Next, we assessed whether this acute depletion before intrasplenic tumor cell injection had an impact on liver metastasis development. To this end, we performed intrasplenic injection of B16OVA cells in mice pre-depleted of macrophages. Macrophage depletion using clodronate resulted in the eventual development of significantly more metastases when analyzed 11 days after intrasplenic injection, while CSF1R blockade alone caused a small apparent increase in metastasis development, which did not reach statistical significance (Figure 2O).

To understand the extent to which effective material engulfing capacities were required to control liver metastasis, we performed an experiment in which mice were subjected to the Nanoprimer strategy, a protocol that aims to saturate engulfing activities of liver macrophages by the injection of a large number of nanoparticles (Saunders, et al. 2020). A single Nanoprimer treatment 2 hours before intrasplenic liver injection increased (close to statistical significance) the development of metastasis in the liver, measured as the area covered by metastatic cells 11 days later (Figure 2P).

Therefore, dynamic liver imaging revealed distinct immune responses upon metastatic seeding. While neutrophils were actively recruited to tumor cells with minimal impact on tumor cell survival, local macrophages exhibited massive engulfment activity that was essential for effective anti-metastatic control.

### Tumor material is retained by local liver macrophages that interact with T cells within the liver tissue

We decided to sequentially explore over time the dynamics of immune cell infiltration and chemokine production in the early steps of liver metastasis. To simultaneously understand the number of myeloid cells infiltrating the tumors and their ability to uptake and retain tumor cell material, mCherry-expressing MC38 tumor cells were intrasplenically injected, and livers were then analyzed at various timepoints: immediately after injection (0 h), and at 4– and 24-hours post-inoculation. Myeloid cell recruitment peaked at 24 hours for most immune cells, except for neutrophils, which peaked at 4h, in line with our observations in intravital imaging experiments (Figure 3A).

**Figure 3.**
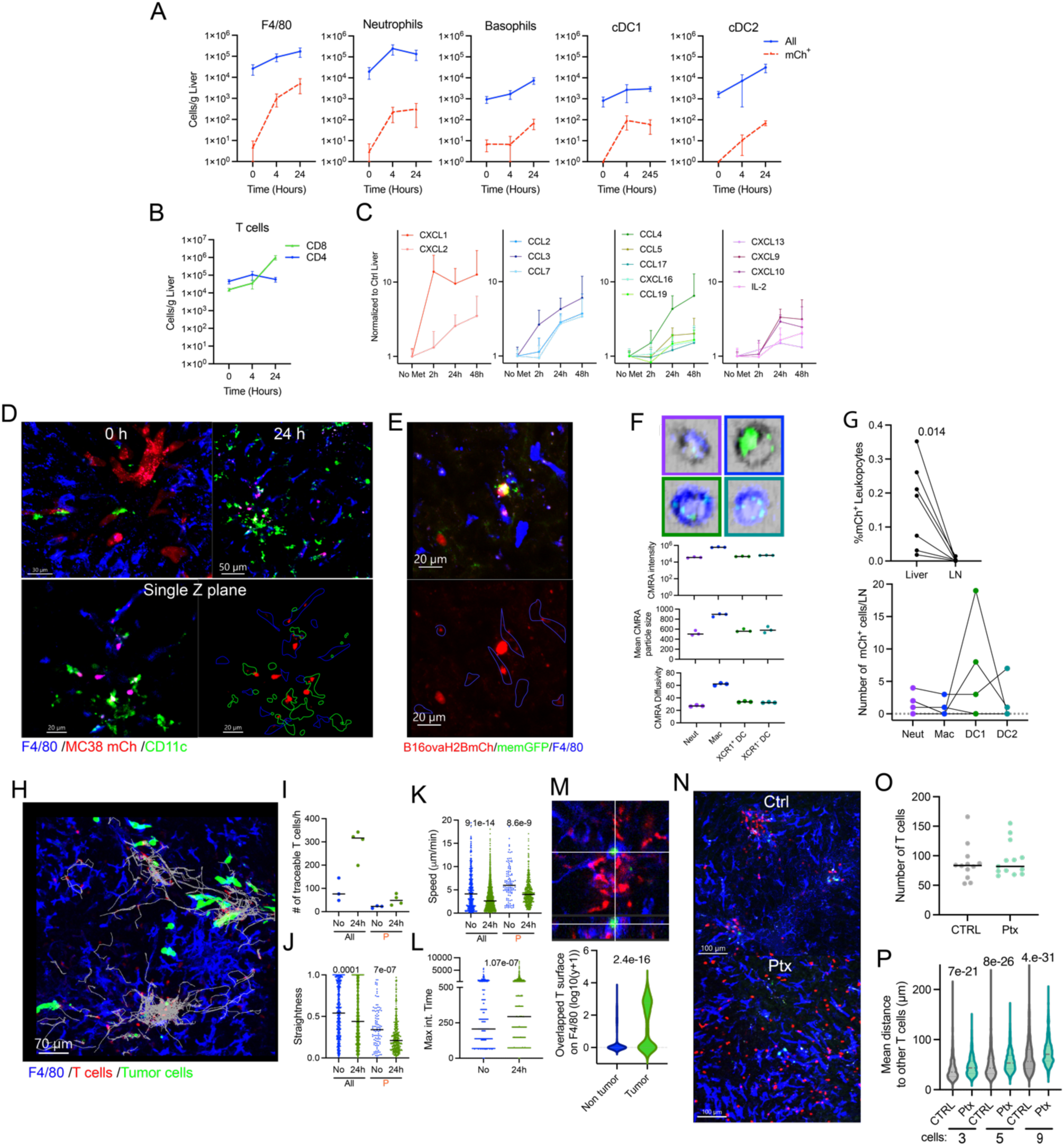
Local liver macrophages retain metastatic tumor cell material and form clusters with locally retained T cells. (A) Flow cytometry quantification of total and tumor material retaining (mCh^+^) myeloid cells in the liver at different time points post-metastasis elicitation. n=6 mice/group. (B) Flow cytometry quantification of T cells in the liver at different time points post metastasis elicitation was assessed. n=6 mice/group. (C) Chemokine profile of the liver at different time points post metastasis elicitation assessed by Luminex. n=6 mice/group. (D) Representative confocal images of MC38 tumor cell material in the liver at the indicated time points. 3D reconstructions are shown above. A single Z-stack of the 24-hour time point, accompanied by a photo of the mCherry channel alone with the shapes of the CD11c and F4/80 positive cells, is shown. (E) Representative confocal images of tumor cell material in the liver at the 24-hour time point post i.s. injection of B16OVA cells expressing GFP in the membrane and mCherry in the nuclei. Single Z plane and single mCherry channel plus F4/80 shape. (F) Imaging flow cytometry of myeloid populations at the 24-hour time point after CMRA-loaded MC38 tumor cell i.s. injection. Representative images and quantification of image-based parameters in each specific population. (G) Flow cytometry assessment of mCherry-positive cells within the liver-draining LNs at 24 hours post i.s. injection time point as compared to the liver tissue (above) and within discrete myeloid populations (below). (H) Representative image of T cell intravital microscopy 24h after metastatic cell arrival. Individual tracks of T cells are shown in white. (I) Quantification of all T cells and patrolling (P) T cells in the livers, in non-metastatic liver and metastatic livers 24h post-metastasis cell injection. Motility parameters of T cells and patrolling T cells (P) in non-metastatic vs 24h post-liver metastasis elicitation. Straightness (J) and speed (K) are shown. (L) Maximum F4/80/ T cell interaction time in non-metastatic and metastatic settings. (M) Representative image of T cells interacting and clustering around macrophages engulfing tumor material (above) and comparative overlapping volume of T cells with macrophages engulfing vs macrophages not engulfing tumor material in the same intravital microscopy videos (below). (N) Representative images of T cells at the 24h post metastasis elicitation time point in mice treated with Ptx after intrasplenic tumor cell injection. (O) Number of T cells quantified in Ptx vs control intravital microscopy videos. (P) Assessment of T-cell clustering based on nearest neighbor distances (3, 5, or 9 neighbors) in control and Ptx-treated conditions. Each dot represents a single cell, except in I and O, where it represents an analyzed video, and in F and G, in which each dot represents a single mouse (for microscopy images, three FOV per mouse analyzed). (H-O) Data were analyzed from at least three videos from two different mice per condition. Medians are represented as a horizontal bar for each condition. Exact p-values are shown for statistical comparison, U Mann-Whitney tests, except for G, in which a paired Student’s T test was used for statistical comparisons.

T cell retention could be observed from the early 4-hour time point, with a pronounced increase in the CD8^+^ subset cell numbers at 24 hours (Figure 3B). In line with these results, multiplex chemokine assessment of the liver tissue revealed two waves of chemokine production. At an early time point, four hours after tumor cell arrival, a strong peak of the neutrophil chemoattractant CXCL1, accompanied by minor increases of monocyte/macrophage/DC chemoattractants CCL7 and CCL4, was observed. A second wave of upregulation of chemokines peaking at 24h and maintained over the next 24 hours included a wide range of myeloid and DC chemoattractants, T and NK cell chemoattractants (CXCL9 and CXCL10) and the IL-2 cytokine, consistent with the active immune cell recruitment observed 24 hours after tumor cell arrival (Figure 3C).

Antigenic material uptake assessed by mCherry inside myeloid cells revealed that all the studied myeloid populations were able to uptake fluorescent material from tumor cells. Already at the 4-hour time point, a fraction of macrophages, neutrophils, and cDCs were positive for tumor cell material, with their numbers usually increasing over time, while basophils only had mCherry^+^ tumor-derived cell material at the later 24-hour time point. Proportionally, macrophages represent the majority of mCherry-positive myeloid cells (>10 fold over other populations), followed by neutrophils and cDC1 at all time points tested (Figure 3A). Three sub-populations were evident within the F4/80^+^ macrophages in our flow cytometry experiments: one characterized by higher levels of F4/80 and medium-to-high levels of MHCII, consistent with resident local macrophages, and two subpopulations within the F4/80^lo^ population with either high or medium-to-low MHCII expression (Figure S3A). The proportion of these populations varied between individual mice, but overall proportions remained similar. However, the proportion of mCherry-positive cells strongly varied between such macrophage populations. A marginal 0.05 % of all the F4/80^lo^ MHCII^lo^ cells showed mCherry positivity, while F4/80^lo^ MHCII^Hi^ and F4/80^Hi^ MHCII^Med^ tend to have 10 and 100-fold more mCherry positive cells, respectively (Figure S3A). These data indicate that tissue resident macrophages are the phagocytic cells with enhanced ability to uptake or retain tumor material. At this time point, liver imaging also showed mCherry-positive macrophages and CD11c EGFP-positive dendritic cells within these two leukocyte populations. mCherry positivity was more pronounced in F4/80 positive macrophages (Figure 3D). Using another cell line, including both surface and nucleus-localized fluorescent proteins (B16OVA PUFFIN mCh), we observed that this tumor engulfment could, in fact, also include nuclear material (Figure 3E). Imaging flow cytometry analysis of tumor cell uptake by myeloid cells revealed that macrophages retained more tumor-derived material, usually stored in large intracellular deposits, which could also be observed in confocal microscopy experiments (Figure 3F). Of note, assessment of mCherry-positive myeloid cells in liver-draining LNs revealed almost no positive leukocytes containing mCherry material, in contrast to the liver tissue. Only a small fraction of cDC1s were positive for mCherry, but in very small numbers and not in all mice (Figure 3G). All these data support that a significant amount of tumor cell material is taken up and retained in the metastatic liver, mainly by local macrophages but also by several other immune cell populations.

We suspected that this tumor cell uptake could enable local macrophages to present or cross-present antigen, as has been described in other pathologies (Muntjewerff, et al. 2020; Venzin, et al. 2025). To check this, we first performed intravital imaging of endogenous T cells, macrophages, and tumor cells in the livers of mice at the 24-hour time point when CD8^+^ T cell numbers peaked. Intravital live imaging evidenced major changes in T cell behavior when compared to T cells within the non-metastatic liver or hours post-tumor cell arrival (Figure 3H). A higher number of T cells was readily observed in the livers, both within the stopping and patrolling classifications (Figure 3I). Frequent clusters of T cells around small tumor particles, many of them engulfed in macrophages, were observed. T cells within such clusters showed evident confined movement characterized by diminished directionality and slower speeds (Figures 3J and 3K). In line with these observations, T cell interactions were more frequent and prolonged with macrophages containing tumor cell fluorescent material (Figure 3L and M). A deeper characterization of T cell clusters using Foxp3GFPxCD2RFP mice revealed the active presence and enrichment of T regulatory cells in these clusters (Figure S3B), indirectly suggesting the formation of antigen-presenting clusters (Natsuaki, et al. 2014). To understand if these T cell clusters were regulated by effective chemotactic T cell recruitment, we performed imaging experiments in mice pretreated with Ptx and found that the presence of such clusters was evidently impaired, although the numbers of T cells within the liver areas retaining tumor cell material remained similar (Figure 3N-P).

A closer look at the videos obtained in T cell reporter mice at 24-hours post-intrasplenic injection showed the appearance of areas enriched in tumor cells, characterized by no F4/80 staining (suggestive of poor blood flow). These areas were also enriched in T cells but with few apparent contacts between T cells and tumor cells. T cells in these non-irrigated, tumor-enriched areas showed different T cell dynamic behavior as compared with T cells in clusters or in areas devoid of T cell enrichment, defined by a strong impairment in motility (Figure S3C and S3D). We hypothesized that this impaired movement could be a result of local hypoxia (Zal and Chodaczek 2010). We therefore used a carbonic anhydrase fluorescent substrate to assess local hypoxia, demonstrating that those F4/80-negative areas were hypoxic and contained tumor cells, unlike the previously described T cell clusters that were negative for carbonic anhydrase activity (Figure S3E and S3F). Cell tracking in hypoxic vs normoxic areas revealed T-cell motility impairment in hypoxic areas (Figure S3G and S3H). Neutrophils in Catchup mice also accumulated in those hypoxic, F4/80-negative regions with apparent motility arrest behavior (Figure S3I)

Overall, our experiments show evidence for adaptive T cell responses in the metastatic liver, defined by the formation of T cell clusters, suggestive of antigen presentation, and the generation of large hypoxic areas enriched in tumor cells that also result in T-cell accumulation and impaired T-cell dynamic behavior.

### Tumor engulfing myeloid cells recruit and activate T cells

To better understand the types of immune cells taking up and retaining tumor material in the liver metastatic site and the effect of such antigen uptake on their function, we performed a scRNAseq on sorted mCherry-positive and negative myeloid cells (CD3, NK1.1, B220 negative; MHC II, CD11b, F4/80 positive) from livers 24 hours after intrasplenic tumor cell injection. We characterized the myeloid populations based on the previously published Liver Cell Atlas (Guilliams, et al. 2022) and identified 16 different immune cell populations present in the metastatic niche, including several monocyte and macrophage populations, several subsets of dendritic cells, basophils, neutrophils, Kupffer cells and lipid-associated macrophages present in the metastatic niche (Figure 4A, 4B and S4A). Cell abundance estimations highlighted that the myeloid composition was dominated by inflammatory monocytes and macrophages, neutrophils, and, to some extent, dendritic cells (Figure 4C). When we focused on the mCherry –positive cells, the cellular landscape was still dominated by inflammatory macrophages, followed by basophils, neutrophils, and Kupffer cells. While highly present in the mCherry negative fraction, cells with limited phagocytic capacity, like monocytes, were much less represented within the mCherry positive cells. When enrichment in populations was defined by comparing mCherry-positive with mCherry-negative cells, basophils were surprisingly the most enriched population, followed by different macrophage subsets, including different subtypes of inflammatory macrophages and Kupffer cells (Figure 4C). Most of our results in the scRNAseq correlated with what we observed in flow cytometry experiments, except for DCs, in which the recovered mCherry-positive cells were fewer than expected based on our previous flow cytometry experiments.

**Figure 4.**
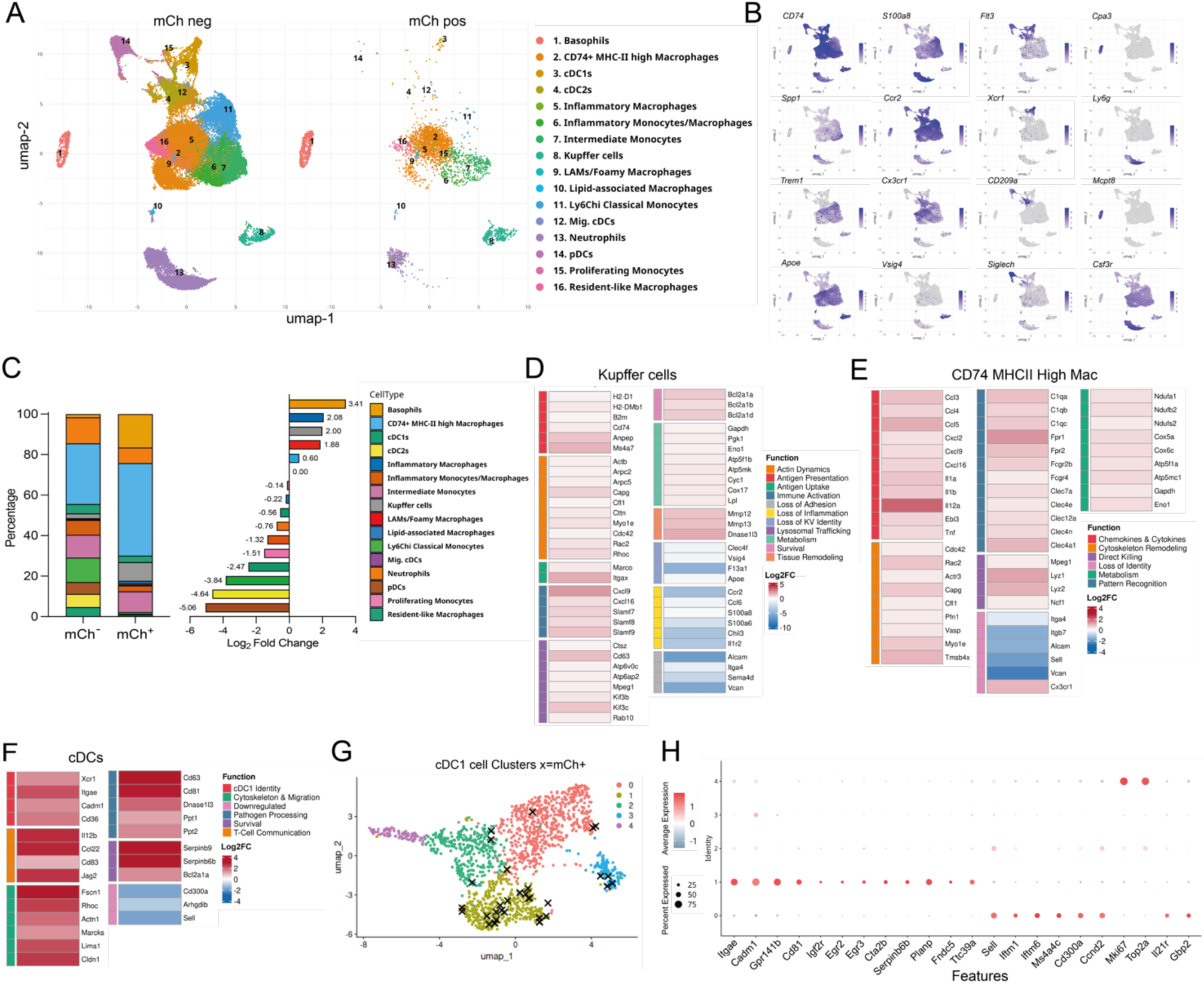
Distinct transcriptional programs of tumor-engulfing myeloid cells. (A) UMAP dimensional reduction of scRNAseq on sorted mCherry^+^ and mCherry^-^ myeloid cells from the liver 24h post metastasis elicitation, colored by cell type. (B) UMAPs depicting myeloid identity-related gene expression. (C) Bar graphs indicating the overall percentage of each myeloid population within each group and the enrichment of each of these populations in mCherry^+^ cells as compared to mCherry^-^. (D) Heatmap of selected differentially expressed genes in mCherry^+^ vs mCherry^-^ in Kupffer cells. (E) Heatmap of selected differentially expressed genes in mCherry^+^ vs mCherry^-^ in CD74^Hi^ macrophages. (F) Heatmap of selected differentially expressed genes in mCherry^+^ vs mCherry^-^ in cDCs. (G) UMAP of re-clustered cDC1s colored by subcluster identification. mCh^+^ individual cDCs1 are depicted with an x overlaid on the graph. (H) Dotplot depicting gene expression analysis on differentially expressed genes identifying cDC subclusters. Data from four pooled mice.

We then investigated specific transcriptional programs elicited by tumor cell uptake in the most relevant isolated immune cell populations. When focusing on Kupffer cells as bona fide resident macrophages in the liver, we observed that mCherry-positive cells showed several relevant processes upregulated when compared to mCherry-negative Kupffer cells (Figure S4B). Notably, a wide number of genes related to antigen presentation were more abundantly expressed in mCherry-positive KCs, including the Ms4a7 gene (Figure 4D and S4B) previously linked to antigen cross-presentation in cDC1s (Xie, et al. 2025). Other major changes involved an increase in genes associated with cytoskeleton dynamics, activation of metabolism and pro-survival programs, along with downregulation of genes related to adhesion and Kupffer cell identity. Together, these changes are consistent with a shift towards a more antigen-presenting phenotype. Importantly, the CXCL16 and CXCL9 T cell chemoattractants were highly upregulated in Kupffer cells containing tumor cell material, suggesting a role for these cells in T cell recruitment and activation after antigen uptake. Other macrophage populations also experienced major transcriptional changes associated with tumor material uptake (Figures S4C and D), including shared functions related to cytoskeleton dynamics and loss of adhesion (Figure 4E) (Acha-Sagredo, et al. 2025). No evident upregulation of antigen presentation-related molecules was observed in these cells, but several proinflammatory cytokines with pro– and antitumor known functions were increased, making it more difficult to interpret the potential role of these antigen-engulfing non-resident macrophages in the antimetastatic immune cell response.

When comparing DEGs in mCherry-positive versus mCherry-negative DCs, we also observed a number of relevant genes upregulated in mCherry-loaded DCs (Figure S4F). Genes associated with survival, cytoskeleton remodeling, and enhanced maturation (CD83, IL12b or DNAse1l3) were clearly upregulated (Fig 4F). Importantly, some genes related to cDC1 identity, like Xcr1 and the Itgae gene encoding for the CD103 integrin, were upregulated, suggesting an enrichment of cDC1s in the antigen-positive population. Reclustering cDC1s in the liver revealed 5 different functional states (Figure 4G and H). Cell cluster 0 was characterized by L-Selectin expression consistent with cells patrolling the liver but not yet activated, cluster 1 was characterized by CD103 expression and markers suggestive of maturation consistent with resident cDC1s or cells undergoing effective maturation. Cluster 4 was characterized as being in a proliferative state, and clusters 2 and 3 shared identity with clusters 4, 0 and 1, respectively, indicating that those may be transitional states. mCherry-positive cDC1s were almost exclusively present in subpopulations 1 and 3, which suggested the initiation of a maturation transcriptional program initiated by antigen uptake (Maier, et al. 2020), indicating that those cells would also be able to effectively cross-present antigen in the liver tissue.

An overlap analysis of genes upregulated in macrophages, Kupffer cells, and dendritic cells allowed us to identify a few common programs upregulated within tumor cell material-positive myeloid cells. Of note CXCL9 was upregulated in both KCs and DCs. All myeloid cells shared a common program of upregulation characterized by cell protection (Serpinb9, Dnase1l3) and vesicle trafficking (Marcks, Cd63) (Figure S4G).

Neutrophils positive for mCherry clustered differently from the whole neutrophil population (Figure 4A). Subclustering of neutrophils based on a recently published transcriptional state atlas (Cerezo-Wallis, et al. 2026) revealed that mCherry-positive neutrophils were composed of both immunosuppressive and antigen-presenting populations (Figure S4H and I) (Bolli, et al. 2026; Yao, et al. 2025), suggesting that antigen-engulfing neutrophils may have dual roles in anti-metastatic liver immune responses and the ability to directly present antigens to T cells in the liver.

Overall, myeloid cells responding to early metastatic infiltration combined immunosuppressive and T cell activation programs, and revealed that resident KCs and cDC1 cells are prone to induce T cell activation during early metastasis immunosurveillance in the liver.

### T-cell reactivation within the metastatic liver is elicited by antigen cross-presentation

Based on our scRNAseq results, we could hypothesize that KCs develop antigen presentation capacities upon tumor material engulfment, making them able to mount local antigen-presenting hubs for further T-cell activation against nascent metastasis.

To understand if proper antigen presentation was behind the formation of T-cell clusters and interactions with macrophages within the liver, we performed intravital imaging of T-cell responses in metastatic livers 24 hours after tumor cell arrival in mice in which we had or had not implanted a previous subcutaneous MC38 tumor. In the absence of a previously implanted tumor, few T cells would likely recognize tumor antigens, making it difficult to mount antigen-specific responses. Image analysis clearly revealed that although patrolling behavior was observed in mice without subcutaneous tumors, the formation of T cell clusters was completely abolished. These results highlight the need for preexisting tumor-specific immunity for effective T cell activation during this early response and demonstrate that this T cell recruitment was not merely guided by the inflammatory response resulting from tumor cell uptake (Figure 5A S13). To model such a response in an antigen-specific model, we adoptively transferred TCR transgenic OTI CD8^+^ T cells recognizing the OVA-derived SIINFEKL peptide in mice with subcutaneous B16OVA tumors. In this model, stronger and more frequent interactions between OT-I cells and local macrophages occurred following intrasplenic injection of OVA-expressing B16 tumor cells compared with OVA-negative B16 cells (Figures S5A-C). As mCherry tracing revealed infrequent tumor cell material arriving at the liver-draining LNs, we decided to compare both local liver and liver-draining LN T-cell activation after intrasplenic tumor injection. Of note, upregulation of activation T-cell markers was evident only within the liver tissue, but not in the liver-draining LNs, indicating that immune surveillance of the liver to metastatic cells was mainly elicited locally (Figure 5B).

**Figure 5.**
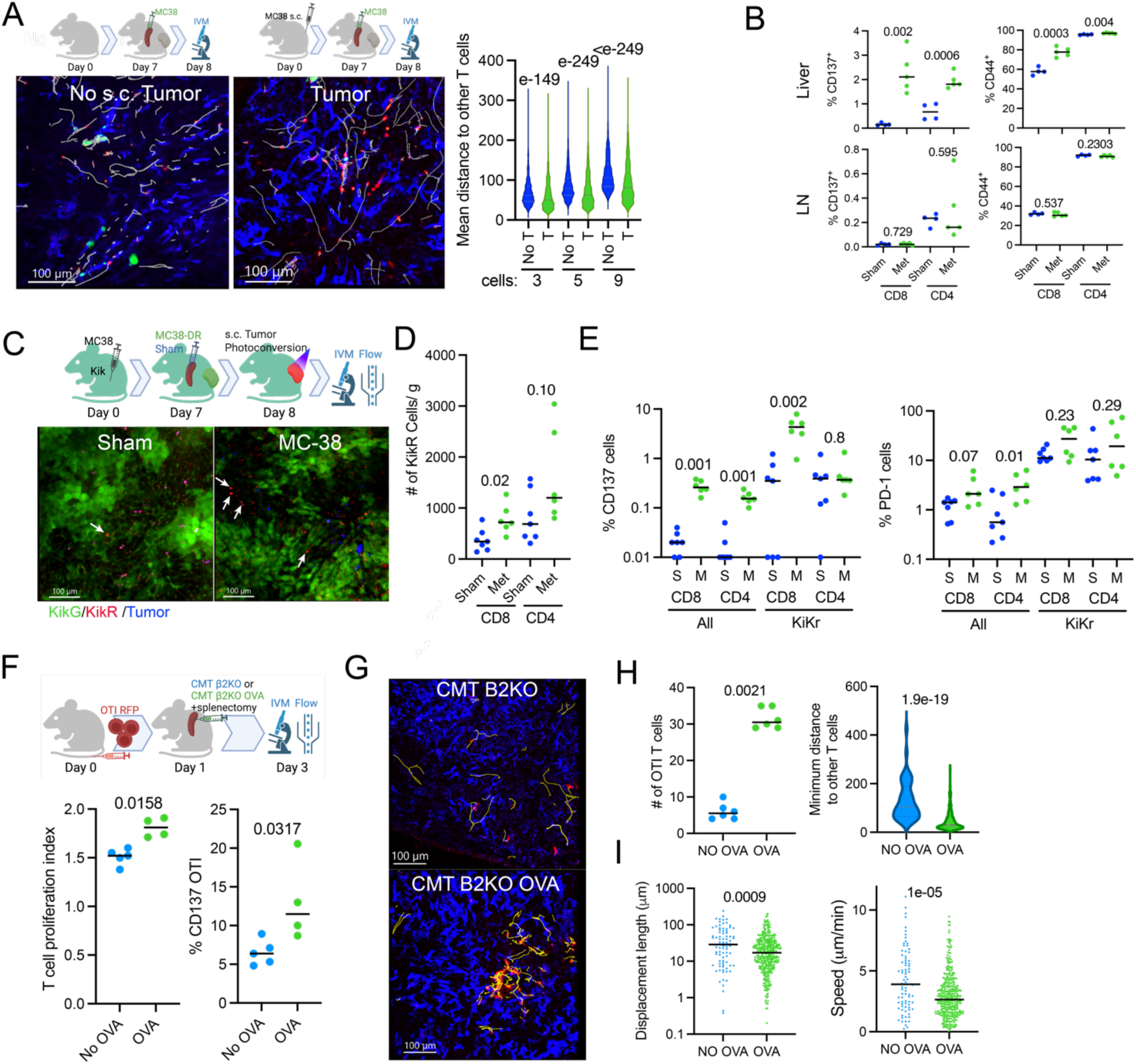
The liver antimetastatic response is characterized by local T-cell reactivation by antigen cross-presentation. (A) Experimental scheme and representative intravital microscopy images depicting T-cell tracks in white, and T-cell-T-cell distance analysis in mice with preexisting subcutaneous MC38 tumors (T) compared with mice without preexisting s.c. tumors (no T), 24 hours after i.s. tumor cell injection. (B) Expression of the indicated T cell activation markers by flow cytometry analysis in the liver and liver-draining LNs 24h post MC38-based metastasis elicitation. (C) Experimental scheme of tumor T-cell tracing experiments in Kikume mice and representative images of the mice’s liver at the indicated time point, with or without metastasis, showing photoconverted Kikume Red positive cell accumulation. (D) Flow cytometry assessment of photoconverted KikRed T-cell infiltration in the liver 24 hours post metastasis. (E) T cell activation markers on all liver-infiltrating T cells and photoconverted KikR T cells in the liver post-metastasis (M= metastasis, S= Sham). (F) Experimental scheme of in vivo cross-presentation assays and flow cytometry assessment of T cell proliferation (CTV dilution) and activation (CD137 expression) within the liver tissue at the indicated post-metastasis time points. (G) Representative intravital microscopy images of RFP^+^ transferred OTI cells in the liver upon metastasis elicitation with OVA-expressing or OVA-negative β2M KO CMT167 cells. T-cell tracks are shown in yellow. (H) Quantification of traceable OTI cells in the liver in intravital microscopy videos and T-cell-T-cell distance upon metastasis elicitation with OVA-expressing or OVA-negative β2M KO CMT167 cells. (I) Motility parameters of fluorescent OTI T cells tracked in the liver upon metastasis elicitation with OVA-expressing or OVA-negative β2M KO CMT167 cells. Data were analyzed from at least three videos from two different mice per condition in imaging experiments. Each dot represents a mouse, except for the left plot in panel H, where it represents an analyzed video, and in right plot of panel H and the whole of panel I, where each dot represents a single cell. Medians are represented as a horizontal bar for each condition. Exact p-values are shown for statistical comparison, U Mann-Whitney tests.

To determine whether T lymphocytes getting activated in the liver were coming from the preexisting subcutaneous tumor in our models, and were mainly antigen-experienced, as suggested by experiments in Figure 5A, we made use of photoconvertible Kikume mice in which UV irradiation allows the conversion of the Kikume green (KikG) protein into red fluorescent (KikR) (Nowotschin and Hadjantonakis 2009). In this model, therefore, UV irradiation of infiltrating immune cells in the s.c tumors would result in a change from green to red fluorescence and would allow their tracking in other distant organs. We performed metastasis elicitation prior to tumor photoconversion and identified photoconverted red fluorescent cells within the liver by imaging and flow cytometry. The injection of the tumor metastatic cells resulted in higher numbers of KikR T cells, particularly in the case of CD8^+^ T cells (Figures 5C and D), with a more activated profile as compared to non-photoconverted T cells present in the liver (Figure 5E), highlighting that at least a proportion of the T cells activated within the metastatic liver have egressed from the tumor site.

We finally aimed to define whether T-cell responses and T-cell activation were being elicited by antigen cross-presentation rather than from tumor cells’ direct antigen presentation. For this purpose, we made use of the highly metastatic lung adenocarcinoma cell line CMT167 (expressing or not OVA) in which the gene for β2-microglobulin (β2M) had been ablated (Perarnau, et al. 1990). Mice were adoptively transferred with Cell Trace Violet (CTV)-stained-red-fluorescent pre-activated OTI cells to mimic a response in which T cells had already been primed in the primary tumor tissue or tumor-draining lymph nodes. One day later, mice were injected intrasplenically with β-2 Microglobulin KO CMT167 cells with or without OVA expression to perform intravital microscopy and flow cytometry analysis on OTI CD8^+^ T cell activation. CTV dilution analysis and CD137 staining revealed OVA-specific activation of OTI cells, indicating efficient cross-presentation in the liver (Figure 5F). In contrast, there was no such evidence for cross-presentation in the liver-draining LNs (Figures S5D and E). Intravital imaging clearly identified that CD8^+^ T cells were forming clusters and modifying their behavior to interact with local macrophages only when OVA cells were injected (Figure 5G to I).

Our findings indicate that these intrahepatic T cell clusters, which form around macrophages positive for tumor-derived material upon tumor cell arrival, function as hubs of antigen cross-presentation to CD8^+^ T cells.

### Both cDC1 and local macrophages are required to elicit antigen cross-presentation in early liver metastasis

Prompted by our scRNAseq data and evidence for local T cell activation within intrahepatic T cell clusters elicited by antigen cross-presentation, we aimed to dissect the mechanisms and cell populations playing a role in local antigen cross-presentation within the liver. First, we found that blocking egress from lymph nodes with FTY720 had no effect on activated T cell levels in the liver, confirming that CD8^+^ T cells were being activated locally rather than in the liver dLNs (Figure 6A). Neutrophil and CD4 depletion resulted in a minor impairment (not significant) of T-cell activation and proliferation, suggesting non-essential cooperation of these two populations in CD8^+^ T cell activation in the metastatic setting (Figure 6A). Basophil depletion led to a minor, non-significant enhanced activation, indicating that the ability of basophils to uptake tumor material was not impacting antigen cross-presentation of metastatic tumor cells in the liver (Figure S6A and B). Finally, clodronate pre-depletion of liver resident macrophages before intrasplenic injection led to an almost complete abolition of T cell activation, indicating that liver macrophages are an absolute requirement for T cell activation (Figure 6B). Analysis of mCherry-positive cells by imaging and flow cytometry upon clodronate treatment revealed that overall, the capacity to capture tumor-derived material was impaired only when local macrophages were depleted, despite the presence of other APC populations. These other APCs tend to increase their tumor material uptake, suggesting a partial compensatory mechanism insufficient to fully restore the loss of the resident macrophage populations (Figure S6C-D).

**Figure 6.**
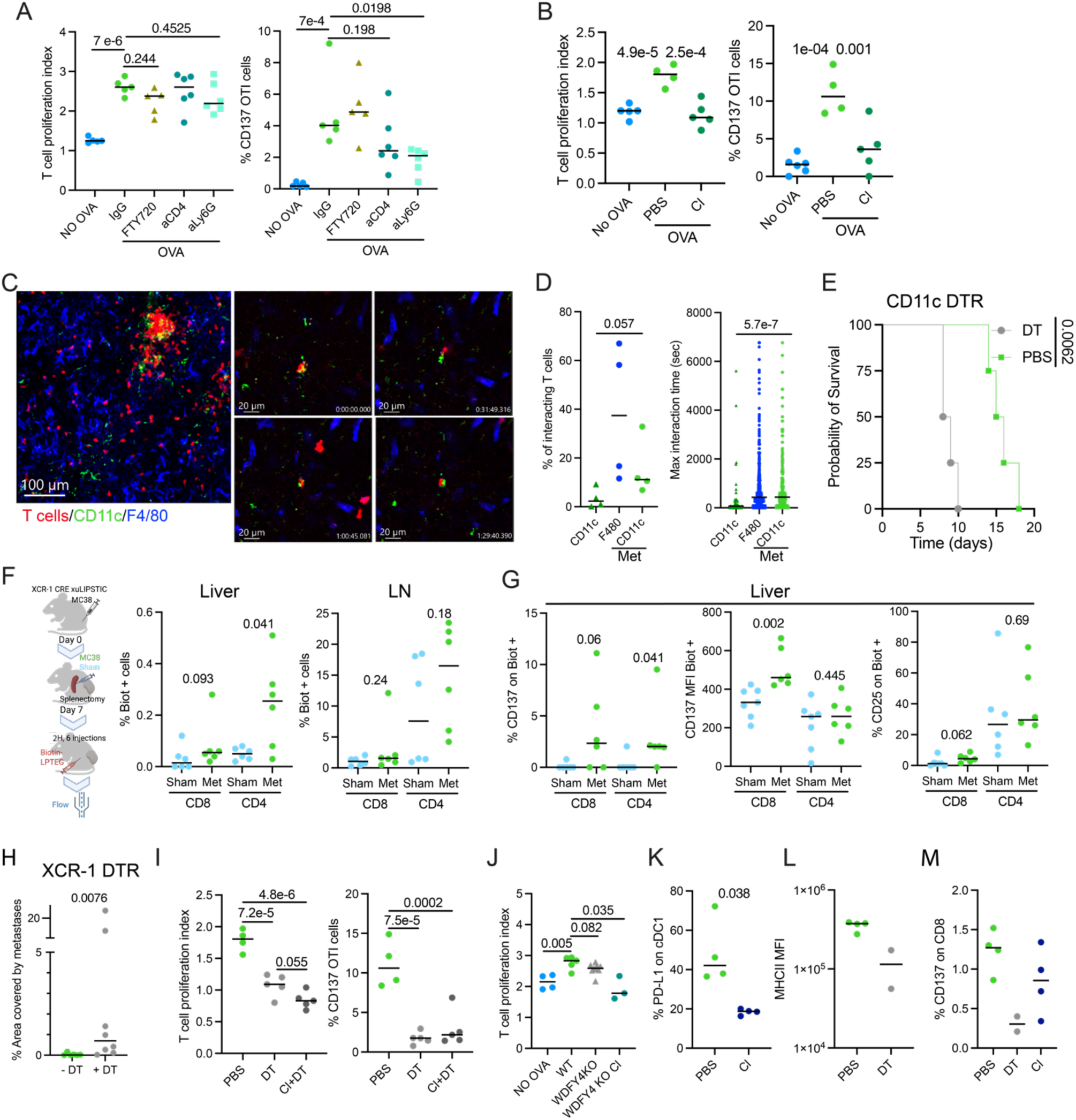
Macrophage and cDC1 cooperation organize anti-metastatic CTL responses. (A) In vivo cross-presentation assays with β2M KO CMT-OVA as in Figure 5F in mice pretreated with the indicated depleting antibodies and pharmacological blockades. Flow cytometry assessment of adoptively transferred OTI cells’ proliferation (CTV dilution) and activation (CD137 surface expression) (B) In vivo cross-presentation assays with β2M KO CMT-OVA as in Figure 5F in mice pre-depleted of macrophages by clodronate liposomes. Flow cytometry assessment of adoptively transferred OTI cells proliferation (CTV dilution) and activation (CD137 surface expression). (C) Representative intravital microscopy images of T-cell responses to liver metastases in CD11c EGFPxCD2RFP double reporter mice showing CD11c cells within T-cell clusters. Sequence of a T cell performing a long-term interaction with a CD11c GFP DC 24h post-metastasis. (D) Quantification of T cells interacting and their maximum interaction times with DCs and macrophages in CD2RFPxCD11cEGFP mice in the premetastatic and 24h post-metastasis time point. (E) Survival assessment of B16OVA bearing CD11cEGFPDTR mice pretreated with DT before i.s. injection of B16OVA. (F) Experimental scheme of cDC1/T cell assessment of interactions using XCR-1uLIPSTIC mice in liver metastasis and flow cytometry assessment of cDC1 contacting (Biotin^+^) T cells within the liver and liver-draining LNs. (G) T cell activation markers on cDC1 contacting T cells in XCR-1uLIPSTIC mice. (H) Quantification of liver metastases in the livers of B16OVA bearing XCR-1-DTR mice pretreated with DT before i.s. injection of B16OVA measured as the area of the liver covered by metastatic cells. (I) In vivo cross-presentation assays with β2M KO CMT-OVA in XCR1-DTR mice pre-depleted of cDC1 by DT injection or both macrophages and cDC1 by DT and clodronate liposomes. Flow cytometry assessment of adoptively transferred OTI cells’ proliferation (CTV dilution) and activation (CD137 surface expression). (J) In vivo cross-presentation assays with CMT-OVA β2M KO in WDFY4^-/-^ cross-presentation-deficient mice pre-depleted (or not) of macrophages by clodronate liposomes. Flow cytometry assessment of adoptively transferred OTI cells’ proliferation (CTV dilution). (K) Expression of PD-L1 maturation marker on liver cDC1 cells by flow cytometry upon clodronate liposomes-mediated macrophage depletion assessed at the 24h post-metastatic time point. (L) Expression of MHC II on liver macrophages by flow cytometry upon DT-mediated depletion of cDC1 in XCR1-DTR mice assessed at the 24h post-metastatic time point. (M) T-cell activation by CD137 expression on liver T cells in mice pre-depleted of cDC1 (DT treatment in XCR1-DTR mice) or macrophages (clodronate liposomes) at 24 hours post-MC38 tumor cell intrasplenic injection. Each dot represents a mouse, except in D, where each dot represents first an analyzed video and then a single cell. Data were analyzed from at least three videos from two different mice per condition in imaging experiments. Medians are represented as a horizontal bar for each condition. p-values are shown for statistical comparison, U Mann-Whitney tests.

Given that dendritic cells were actively retaining tumor cell material within the liver and upregulating antigen-presenting features at the transcriptomic level, we performed intravital live imaging of DC within the metastatic liver. Dendritic cells were frequently observed within liver T cell clusters, performing interactions with T cells and displaying similar interaction dynamics to those observed for macrophages (Figure 6 and D). DC depletion just before metastasis elicitation using a CD11c DTR mouse model, in which a single diphtheria toxin (DT) injection can lead to a transient depletion of dendritic cells, led to shorter survival in these mice, indicating a role for DCs in liver metastasis immunosurveillance (Figure 6E).

Since cDC1s are highly specialized in antigen cross-presentation and represent the predominant intrahepatic DC population retaining tumor antigen, we sought to investigate their role in this metastatic setting. We used the XCR-1^iCRE^x ROSA26^uLIPSTIC^ (XCR-1-uLIPSTIC) cell-cell contact tracing model (Nakandakari-Higa, et al. 2024) to trace cDC1-T cell interactions and the amount and functional state of T cells that had interacted with cDC1s in the liver. XCR1-uLIPSTIC mice were injected intraperitoneally for two hours with the uLIPSTIC labeling reagent LPETG-biotin. T cells were then recovered and the percentage of biotin-positive cells and their activation markers were measured by flow cytometry. Liver metastasis elicitation increased the percentage of biotin-positive T cells, mostly within the liver and not in the liver-draining LNs (Figure 6F). The activation markers CD137 and CD25 were almost absent in biotin-positive T cells in non-metastatic conditions and showed upregulation only in mice with liver metastases, indicating that CD8^+^ T cells interacting with cDC1s in the liver were the ones recognizing antigen (Figure 6G). To define the role of cDC1s in the elicitation of anti-tumor liver T-cell responses, we used XCR1-DTR mice, a cDC1-depletable mouse model. In this setting, cDC1 acute depletion with two doses of DT led to a strong reduction of cDC1 cell numbers in the liver that was partially recovered and completely recovered at days 5 and 8 post-depletion, respectively, restricting depletion to the early phases of metastatic colonization (Figure S6F). To understand the actual relevance of cDC1 in liver metastasis control, we performed acute cDC1 depletion before metastasis elicitation and found that liver metastasis control was fully dependent on cDC1 presence at the initial steps of metastatic colonization (Figure 6H). In this setup, liver antigen cross-presentation to OTI cells was strongly limited, and clodronate-based macrophage depletion, in combination with DT treatment, led to a further decrease in T cell proliferation that approached statistical significance, suggesting some degree of cooperation between both myeloid populations to allow proper T cell reactivation by antigen cross-presentation (Figure 6I).

To distinguish between the roles of antigen cross-presentation and other functions of cDC1s in this setting, we used WDFY4-deficient mice in which cDC1s cannot cross-present cell/antibody-associated antigens and treated them with clodronate. In this mouse model, the T cell proliferation response was partially inhibited (though not significant). However, clodronate treatment completely abolished T cell activation in WDFY4^-/-^ mice, confirming the importance of resident macrophages, and suggesting that cDC1 functions other than cross-presentation may be of greater importance in this local liver activation of T cells (Figure 6J).

Given that both cDC1 and macrophages were required for proper T cell activation, we performed cDC1 and macrophage depletion studies and looked at the endogenous T cell response to MC38 metastatic cells and the relative effects of each myeloid population on the other. Clodronate depletion did not alter the numbers of liver cDC1s (Figure S6G), as previously demonstrated, but it did alter their maturation state as PD-L1 levels were much decreased on cDC1s (Figure 6K). cDC1 depletion did not alter macrophage numbers in the liver either (Figure S6H), but the levels of MHC II expression on macrophages were much lower, suggesting a weaker inflammatory response when cDC1s were absent (Figure 6L). In line with these observations, T cell numbers were not substantially changed within the liver when either one of the populations was depleted (Figure S6I), but the formation of T cell clusters around tumor cell fragments did, although such inhibition was much stronger when depleting cDC1s (Figure S6K). Importantly, T cell activation was only fully inhibited when cDC1s were absent (Figure 6M and S6J), suggesting that cDC1s are indispensable for the elicitation of T cell endogenous responses, at least by favoring the formation of T cell antigen-presenting hubs within the liver. cDC1 and macrophage depletion lead to alterations in chemokines expressed in the metastatic liver. Relevant chemokines, such as CCL19 and CCL5, were less abundant when both populations were absent; other chemokines showed reduction only evident upon cDC1 depletion (CXCL9) or clodronate treatment (CXCL13, CCL4) (Figure S6L), denoting the non-redundant roles of both populations in establishing immune responses in the liver in early metastasis.

All these data suggest effective local macrophage-cDC1 cooperation to elicit T-cell immune surveillance of metastasis within the liver tissue, uncovering an unexpected role for local tissue macrophages in taking up tumor cell material, attracting and promoting T cell responses, but with required cooperation of cDC1 cell recruitment to elicit effective T cell activation and proliferation, and as a consequence, liver metastasis control.

### cDC1 expansion increases liver immunity and metastasis control

To understand if any of the mechanisms of immune surveillance performed by liver antigen-presenting cells could be exploited in cancer immunotherapy, we then used mice bearing B16OVA subcutaneous tumors and treated the mice with either a single dose of Flt3L-Ig to expand cDC numbers or a single dose of anti-CD47 antibodies to facilitate antigen phagocytosis (Tseng, et al. 2013). Anti-CD47 treatment led to no evident control of liver metastases, while Flt3L-Ig almost completely inhibited the formation of liver metastases in mice (Figure 7A). In a second experiment in which splenectomy was performed upon intrasplenic injection of B16OVA cells, cDC1 expansion by Flt3L-Ig again led to an almost complete inhibition of metastasis formation (Figure 7B). In vivo antigen cross-presentation assays revealed that Flt3L-Ig pretreatment greatly enhanced T cell proliferation in the liver (Figure 7C), indicating that Flt3L-based therapies may be of use to boost an effective anti-metastatic immune surveillance within the liver.

**Figure 7.**
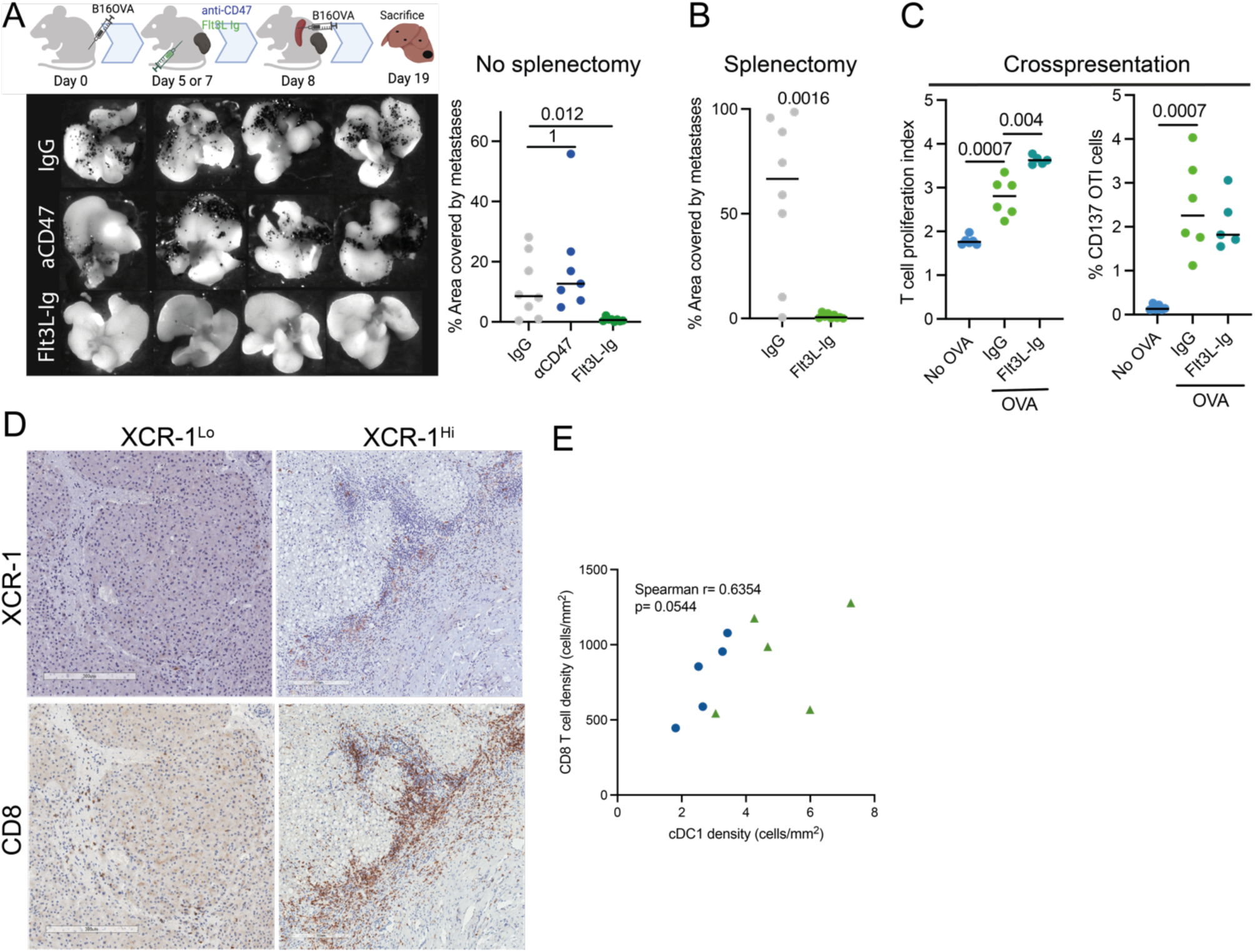
cDC1 expansion allows effective immunosurveillance of liver metastases. (A) Experimental scheme of liver metastasis control experiments in the B16OVA model, in mice pretreated with Flt3L-Fc or anti-CD47 blocking antibodies. Representative images of B16OVA metastases in livers and quantification of liver metastatic surface at the indicated time point of sacrifice. (B) Experiments as in A, but performing splenectomy right after B16OVA intrasplenic injection. (C) In vivo cross-presentation assays after β2M KO CMT167 OVA i.s. injection showing OTI proliferation (CTV dilution) and activation (CD137 surface expression) upon Flt3L-Fc-based DC expansion. (D) Representative IHC of XCR1 and CD8 staining on liver metastases showing two cases with differential XCR1^+^ cell infiltration. (E) Correlation of CD8^+^ and XCR1^+^ cell/ density in two series of liver metastases from colorectal and pancreatic tumors. Each dot represents a mouse/patient. Medians are represented as a horizontal bar for each condition. Exact p-values are shown for statistical comparison, U Mann-Whitney tests, except for E, where a non-parametric Spearman correlation test was performed.

To understand the relevance of cDC1s in liver metastasis immunosurveillance in human cancer patients with liver metastases, we analyzed two small cohorts of colorectal and pancreatic cancer patients with liver metastases and quantified the numbers of cDC1s and CD8^+^ T cells by conventional IHC in serial sections of liver metastases biopsies. Our analysis showed a clear correlation between CD8^+^ T cells and cDC1s (Figure 7D and E), highlighting that CD8^+^ T cell immune surveillance of the liver metastases was connected to cDC1 infiltration as observed in our experimental data in mice.

Therefore, pharmacological cDC1 expansion in cancer patients may enhance immune surveillance and improve metastasis prevention or control of liver metastases.

## DISCUSSION

Immune surveillance mechanisms against metastasis in organs remain largely unknown and understudied in tumor immunology. Most of the studies defining immune microenvironments at the metastatic sites have focused on the characterization of already established metastases, rather than the responses elicited during early dissemination and arrival of malignant cells to metastatic organs. Responses against pioneering metastatic cells may have similar features irrespective of the target organ, but also tissue-specific mechanisms that can define how immune responses are set against evolving metastasis. Differential immune mechanisms may drive immune escape that facilitates metastatic outgrowth and progression in specific target organs.

Using fluorescence tracing, scRNA seq, and intravital imaging, we have observed strong similarities to the multistep cascade elicited by pioneering metastatic cells within the liver as compared to those previously described in the lung. In this regard, the generation and release of large vesicles, which may be induced by flow-induced stress effects, as described by Headley and colleagues (Headley, et al. 2016), was also observed, along with a fast and acute neutrophilic response, in this case characterized by rapid interactions with tumor cells followed by movement away from the target tumor cells. The fact that mCherry fluorescent material was readily observed by flow cytometry in neutrophils early upon tumor cell arrival suggests previously characterized phagocytic or trogocytic behaviors in neutrophils interacting with tumor cells (Belz, et al. 2026; Matlung, et al. 2018). Yet, imaging approaches could not clearly evidence antigen uptake in the observed interactions, unlike for macrophages, which were often observed taking up large amounts of tumor material upon nibbling entire tumor cells or interacting with large flowing tumor cell fractions. The fate of all these neutrophils with engulfed tumor cell material and their ability to mediate first-hand or second-hand tumor antigen presentation remains to be explored (Wu, et al. 2024b). Transcriptomic data revealed that a large proportion of tumor-engulfing neutrophils showed a previously described antigen presentation program. Neutrophil depletion limited antigen cross-presentation to CD8^+^ T cells at least to a certain extent, suggesting that neutrophils may be involved in direct or, most likely, second-hand antigen presentation.

Despite the similarities observed with myeloid recruitment and tumor material uptake in the lungs, our investigations in the liver revealed a clear liver-specific immune response characterized by massive antigen material engulfment, mainly performed by local macrophages (characterized by F4/80 ^high^ expression levels) (Gul, et al. 2014; Kimura, et al. 2016). In addition, the elicitation of T-cell dynamic clusters, characterized by strong and persistent interaction of T cells with macrophages and DCs, was observed within the liver but not within the metastatic lung, in which responses in lung-draining LNs were found to be more relevant. The formation of these local responses within the liver parenchyma has been clearly evidenced during the elicitation of antiviral responses within the liver (Huang, et al. 2013), but our data show that the liver has the capacity to direct and prepare such local immune responses only when concurrent antigen material uptake and proper antigen presentation co-occur. In line with this observation, both local macrophages and cDC1s are equally required for effective cross-presentation of cell-associated antigen and eventual liver metastasis control.

The fact that WDFY4-deficient mice, which have strong defects in antigen cross-presentation by cDC1s (Theisen, et al. 2018), only partially lose the ability to cross-present antigen to TCR-specific anti-OVA cells in our experiments suggests important antigen-presenting roles for other cells. Furthermore, the fact that evident antigen presentation machinery upregulation is specifically upregulated in Kupffer cells suggests that local macrophages can also drive tumor antigen presentation. However, cooperation with cDC1s locally, most likely through IL-12 production (upregulated in antigen-bearing cDC1s), is locally required to drive effective antitumor CTL responses and eventual metastasis control. Kupffer cells have been shown to be able to cross-present viral antigens (Benechet, et al. 2019; Venzin, et al. 2025), but only with help provided by CD4^+^ T cells. In our hands, similarly to WDFY4, CD4^+^ T cell depletion resulted in a mild, not significant impairment of CD8^+^ T activation in cross-presentation studies. CD4 depletion could have limited Treg responses, as Tregs were frequently present in antigen-presenting hubs (Devi-Marulkar, et al. 2022), or the particular strength of OTI TCR may involve no requirement for T cell help in this setting. Of note, our findings strongly reveal that local liver draining LNs have, if any, only a minor contribution in the elicitation of the antimetastatic response at these early time points, although it is true that our experimental systems do not allow us to determine if the spleens may have a more relevant role as an antigen-draining site for the liver antimetastatic immune responses.

An otherwise surprising observation in our experiments was to see that a single dose of Flt3L-Fc protein (Decalf, et al. 2025) before metastatic seeding almost completely impaired the development of liver metastasis. Although Flt3L alone rarely exerts an anti-tumor effect against established tumors, its combination with immunotherapy has been shown to be highly synergistic (Lai, et al. 2026; Salmon, et al. 2016; Sanchez-Paulete, et al. 2016). The fact that cDC1 expansion may favor metastasis immunosurveillance, even here in a context in which an advanced primary tumor was present, may support further development of this therapeutic approach, whose clinical progress has been limited by the appearance of an acute leukemia in a patient with previous clonal hematopoiesis as a result of the treatment with recombinant Flt3L during a phase I trial (Anandasabapathy, et al. 2015).

To be able to visualize and effectively quantify the limited T cell responses against pioneering tumor cells, we have had to rely on a simplified model of liver metastasis, characterized by the injection of a larger number of cells than those expected to reach the tissue. This is an evident limitation of our study, partially compensated for by our imaging approach, identifying local responses in the areas of interest. Furthermore, some features of the metastatic emboli, like platelet and neutrophil enrichment, multicellular tumor cell composition, or specialized metastatic phenotypes of tumor cells, may not be fully recapitulated by our model (Labelle, et al. 2014; Szczerba, et al. 2019; Tauriello, et al. 2018). Despite these limitations, we believe that our work dissects for the first time the spatiotemporal response elicited during antimetastatic surveillance, highlighting the ability of the liver to overcome its tolerogenic state when tumor cells reach the tissue and defining the key role of cDC1 in liver immunosurveillance and organization of anti-tumor responses within liver metastasis, as supported by our data in human specimens.

Overall, our work highlights that the liver has the potential to generate antitumor immune responses to arriving metastatic tumor cells upon their initial colonization steps, combining active tumor material engulfment by myeloid cells, local antigen cross-presentation and myeloid-coordinated T cell activation, which can be further enhanced by therapeutic interventions like dendritic cell expansion to limit metastasis at its earliest stages.

## ACKNOWLEDGMENTS

We thank Diego Alignani (Flow Cytometry Facility); Eneko Elizalde, Alberto Espinal, and Elena Ciordia (Animal Facility) and Carlos Ortiz de Solorzano, Ainhoa Urbiola, Cristina Ederra and Xabier Morales (Light imaging services) for their excellent technical support.

## FIGURES AND FIGURE LEGENDS

**Figure S1.**
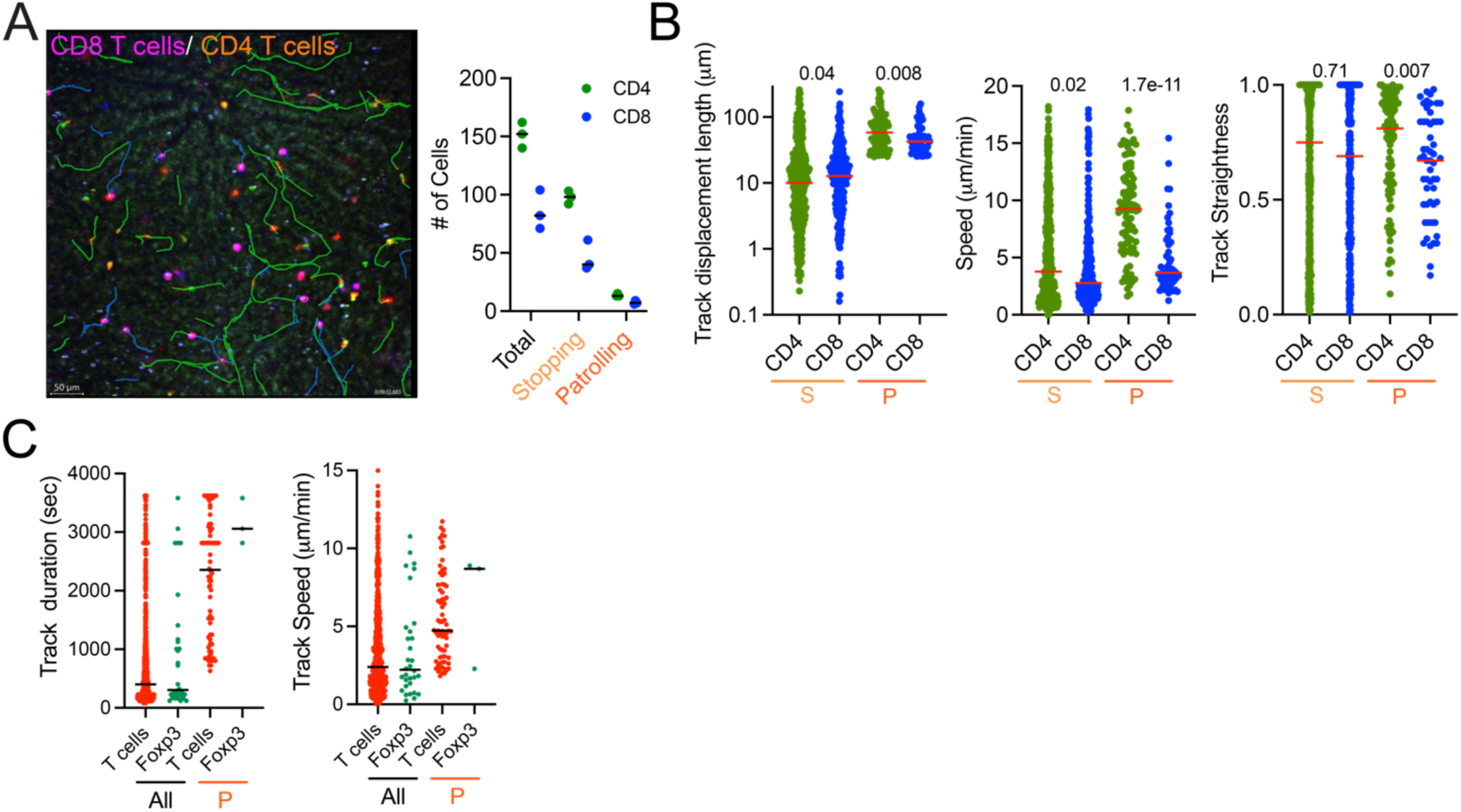
Gi signaling and liver resident macrophages contribute to T-cell patrolling behavior within the liver. Related to Figure 1. (A) Representative image of CD4 (yellow) and CD8 (purple) RFP positive T cells and their tracks (green and blue lines respectively) and quantification of T cells with different motility behaviors. (S= stopping, P=Patrolling) (B) Comparative assessment of motility features of overall and patrolling CD4^+^ and CD8^+^ T cells. (C) Comparative assessment of motility features of overall and patrolling CD2RFP and CD2RFP FOXP3 EGFP T cells. Data were analyzed from at least three videos from two different mice per condition. Each dot represents a single cell, except in A, where it represents an analyzed video. Medians are represented as a horizontal bar for each condition. Exact p-values are shown for statistical comparison, U Mann-Whitney tests.

**Figure S2.**
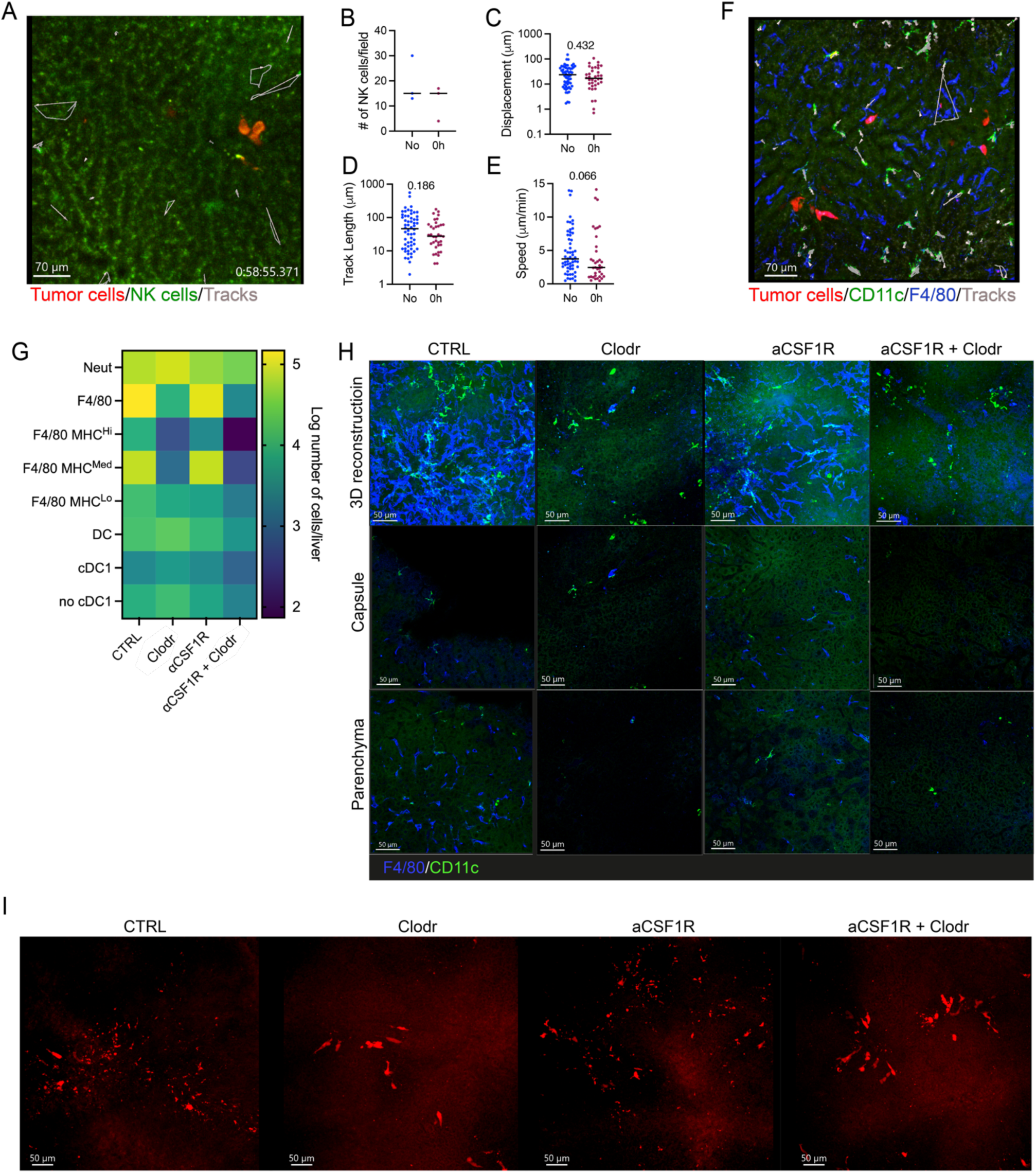
Absence of NK– and DC-directed chemotaxis towards arriving tumor cells and effects of macrophage depletion strategies in the liver. Related to Figure 2. (A) Representative image of NK cell behavior upon metastatic cell arrival. Individual tracks of NK cells are shown in white and grey arrows show the direction of the overall displacement of each tracked cell. (B) Quantification of tracked NK cells in intravital microscopy videos in non-metastatic livers and immediately after intrasplenic injection. Motility parameters, including displacement (C), length (D) and speed (E) of tracked NK cell tracked cells in intravital microscopy videos in non-metastatic livers and immediately after intrasplenic injection. (F) Representative image of NK cell behavior upon metastatic cell arrival. Individual tracks of NK cells are shown in white and grey arrows show the direction of the overall displacement of each tracked cell. (G) Heatmap showing the absolute number of different myeloid populations following 48 h of treatment with α-CSF1R, clodronate liposomes or a combination of both at the 24h time point post i.s. injection. (H) Fluorescent microscopy imaging of clarified liver tissue showing macrophages (F4/80^+^) and DCs (CD11c^+^) following 48 h of treatment with α-CSF1R, clodronate liposomes, or a combination of both at the 24h time point post i.s. injection showing F4/80 macrophages and CD11c DCs. (I) Representative images of mCherry MC38 tumor cells visualized by whole liver confocal imaging tissue following 48 h of treatment with α-CSF1R, clodronate liposomes, or a combination of both at the 24h time point post i.s. injection. (A-F) Data were analyzed from at least three videos from two different mice per condition. Each dot represents a single cell. Medians are represented as a horizontal bar for each condition. p-values are shown for statistical comparison, U Mann-Whitney tests. (G-I) 7-9 mice per group.

**Figure S3.**
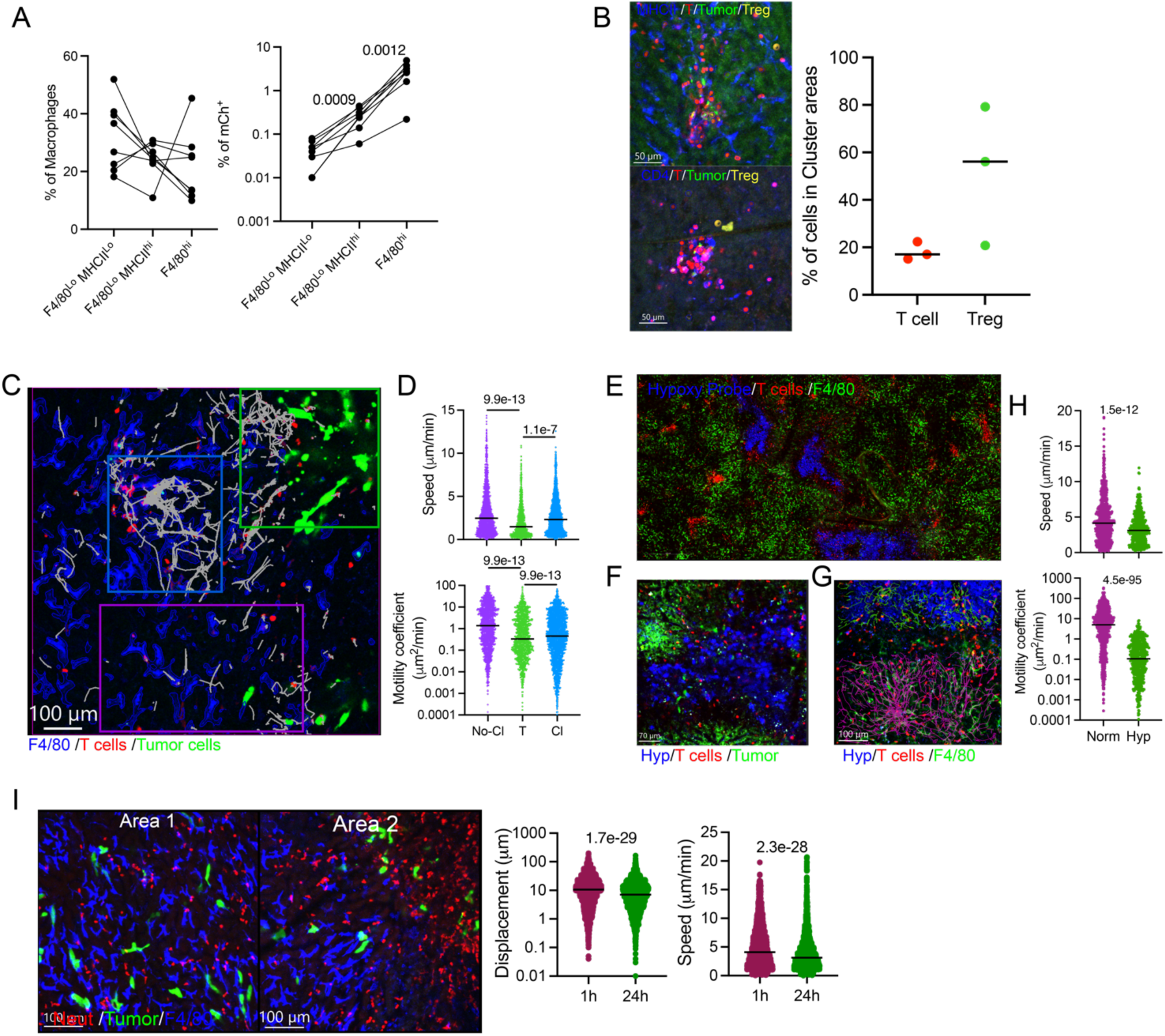
T cells accumulate in hypoxic regions enriched in tumor cell clusters. Related to Figure 3. (A) Subclassification of liver macrophages based on F4/80 and MHC II expression levels. Proportion of all F4/80^+^ macrophages at the 24h post metastasis time point (left) and proportion of tumor material bearing (mCherry positive) cells of each subset. (B) Representative images of T regulatory cells in T cell clusters in intravital videos performed in Foxp3 EGFPxCD2RFP mice and quantification of Treg tracks within clusters and far from clusters. (C) Representative image of an intravital microscopy video performed at the 24h time point post-tumor cell injection, highlighting three regions: normal parenchyma (no-CI), parenchyma enriched in T cell clusters and macrophages containing tumor material (CI), parenchyma enriched in tumor cells and negative for F4/80 macrophages (T). Tracks are shown in white. (D) Comparison of motility parameters of T cells within each type of region. (E) Tile scan of a 24h-post metastasis liver upon injection of the Hypoxia tracker in CD2RFP mice showing large hypoxic regions surrounded by T cells. (F) Representative image of tumor cell clusters within hypoxic regions. (G) Representative image of T cells tracked in intravital imaging videos in hypoxic versus non-hypoxic regions. (H) Motility parameters of T cells within hypoxic vs non-hypoxic regions. (I) Motility parameters of neutrophils in intravital microscopy videos performed in Catchup mice, comparing early vs 24h post metastatic seeding. Representative images show F4/80 negative areas enriched in clustered tumor cells and neutrophils. Data were analyzed from at least three videos from two different mice per condition. Each dot represents a single cell, except in B, where it represents an analyzed video, and in A, where each dot represents a single mouse. Medians are represented as a horizontal bar for each condition. p-values are shown for statistical comparison, U Mann-Whitney tests

**Figure S4.**
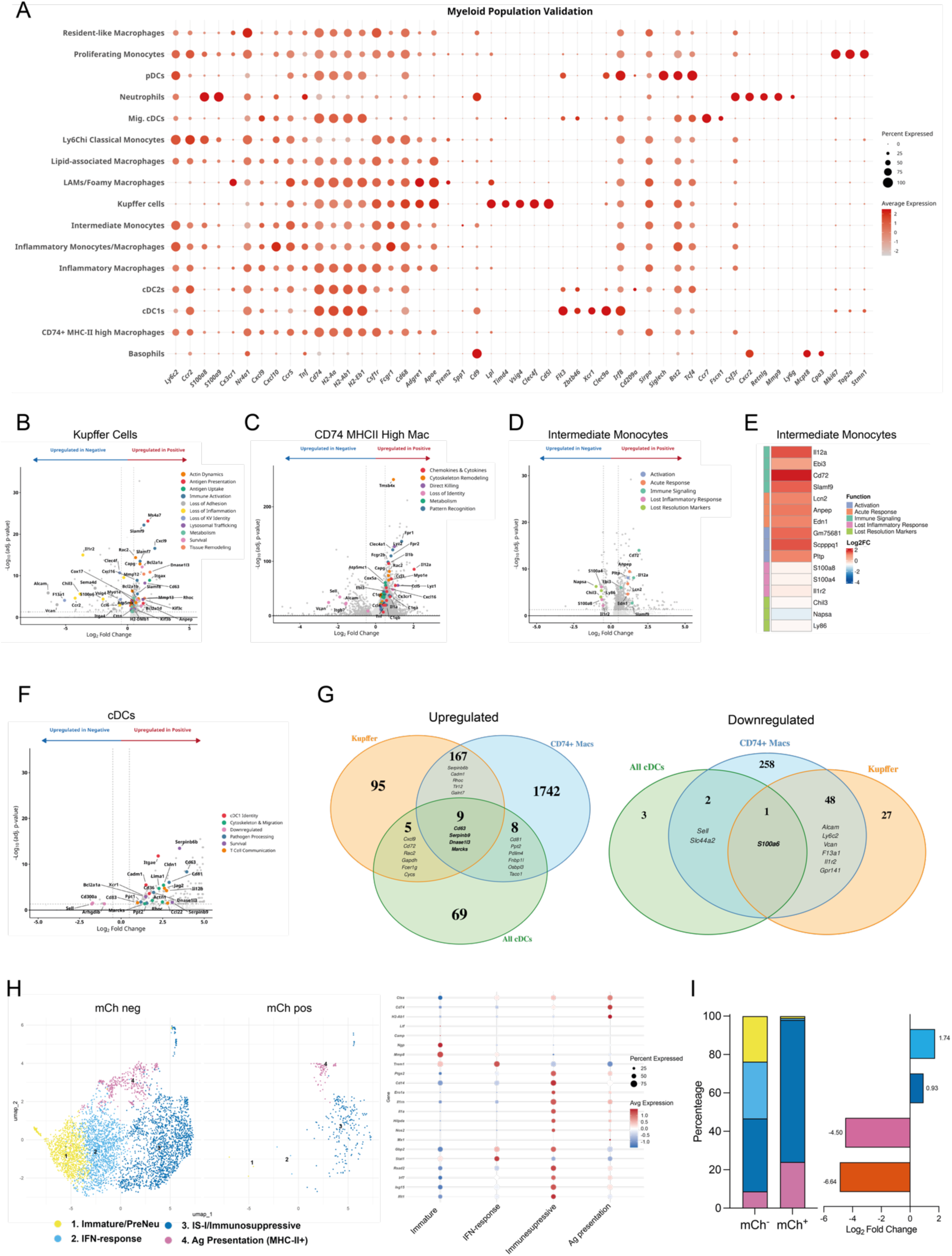
scRNAseq on tumor-material-bearing myeloid cells in nascent liver metastasis. Related to Figure 4. (A) Dotplot depicting gene expression analysis on differentially expressed genes identifying liver myeloid cells. (B) Volcano plot showing differentially expressed genes comparing mCh^+^ and mCh^-^ Kupffer cells and biological process-based colour-coded selected genes. (C) Volcano plot showing differentially expressed genes comparing mCh^+^ and mCh^-^ CD74^Hi^ Macrophages and biological process-based colour-coded selected genes. (D) Volcano plot showing differentially expressed genes comparing mCh^+^ and mCh^-^ Intermediate Monocytes and biological process-based colour-coded selected genes. (E) Heatmap showing selected differentially expressed genes in mCherry^+^ vs mCherry^-^ in Intermediate Monocytes. (F) Volcano plot showing differentially expressed genes comparing mCh^+^ and mCh^-^ cDCs and biological process-based colour-coded selected genes. (G) Venn diagram showing the overlapping of upregulated vs downregulated genes in tumor material positive Kupffer cells, CD74^hi^ Macrophages and cDCs. (H) UMAP of re-clustered neutrophils according to the NeuMap system, colored by subcluster identification and Dotplot depicting gene expression analysis on differentially expressed genes, identifying neutrophil subclusters. (I) Bar graphs indicating the overall percentage of each neutrophil population within each group and the enrichment of each of these populations in mCherry^+^ as compared to mCherry^-^ neutrophils. Data from 4 pooled mice.

**Figure S5.**
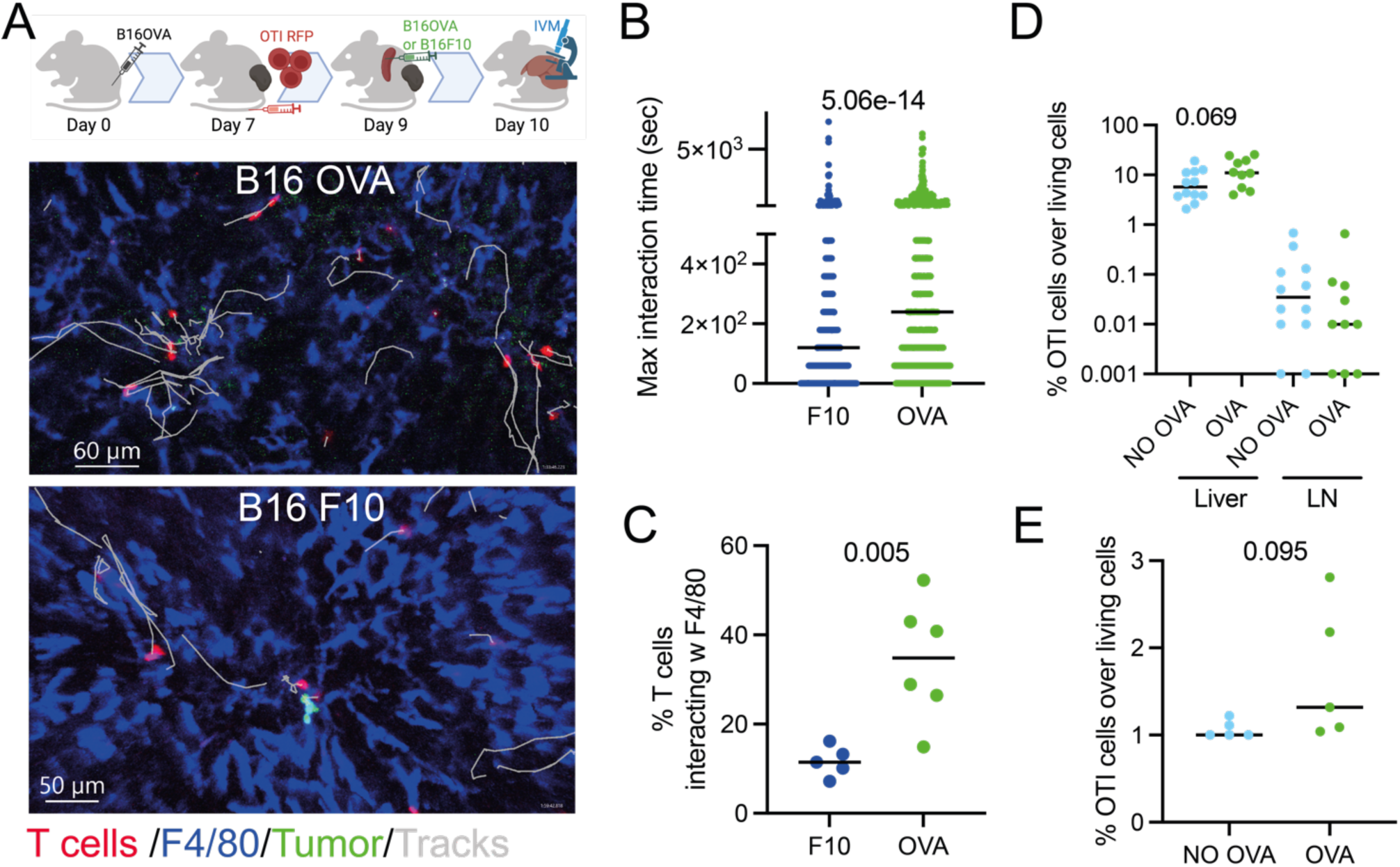
Antigen-specific T-cell responses are elicited within the liver tissue, but not within the liver-draining LNs in liver metastasis. Related to Figure 5. (A) Experimental scheme and representative intravital microscopy images, depicting T cell tracks in white, of antigen-specific T cell responses in the liver upon fluorescent OTI T-cell injection in mice bearing s.c. B16OVA after liver metastasis elicitation with B16OVA or B16F10 tumor cells. (B) Max interaction time of tracked T cells in the liver with F4/80 macrophages in both conditions. (C) Percentage of T cells engaging with macrophages in both conditions. (D) Quantification of OTI infiltration in the liver and liver-draining LNs in cross-presentation assays, as in Figure 5F. (E) OTI proliferation measured as CTV dilution in liver-draining LNs upon metastasis elicitation with OVA-expressing or OVA-negative β2M KO CMT167 cells. Data were analyzed from at least three videos from two different mice per condition in imaging experiments. (B) Each dot represents a cell, (C) each dot represents a video, (D) and (E) each dot represents a mouse. Medians are represented as a horizontal bar for each condition. Exact p-values are shown for statistical comparison, U Mann-Whitney tests.

**Figure S6.**
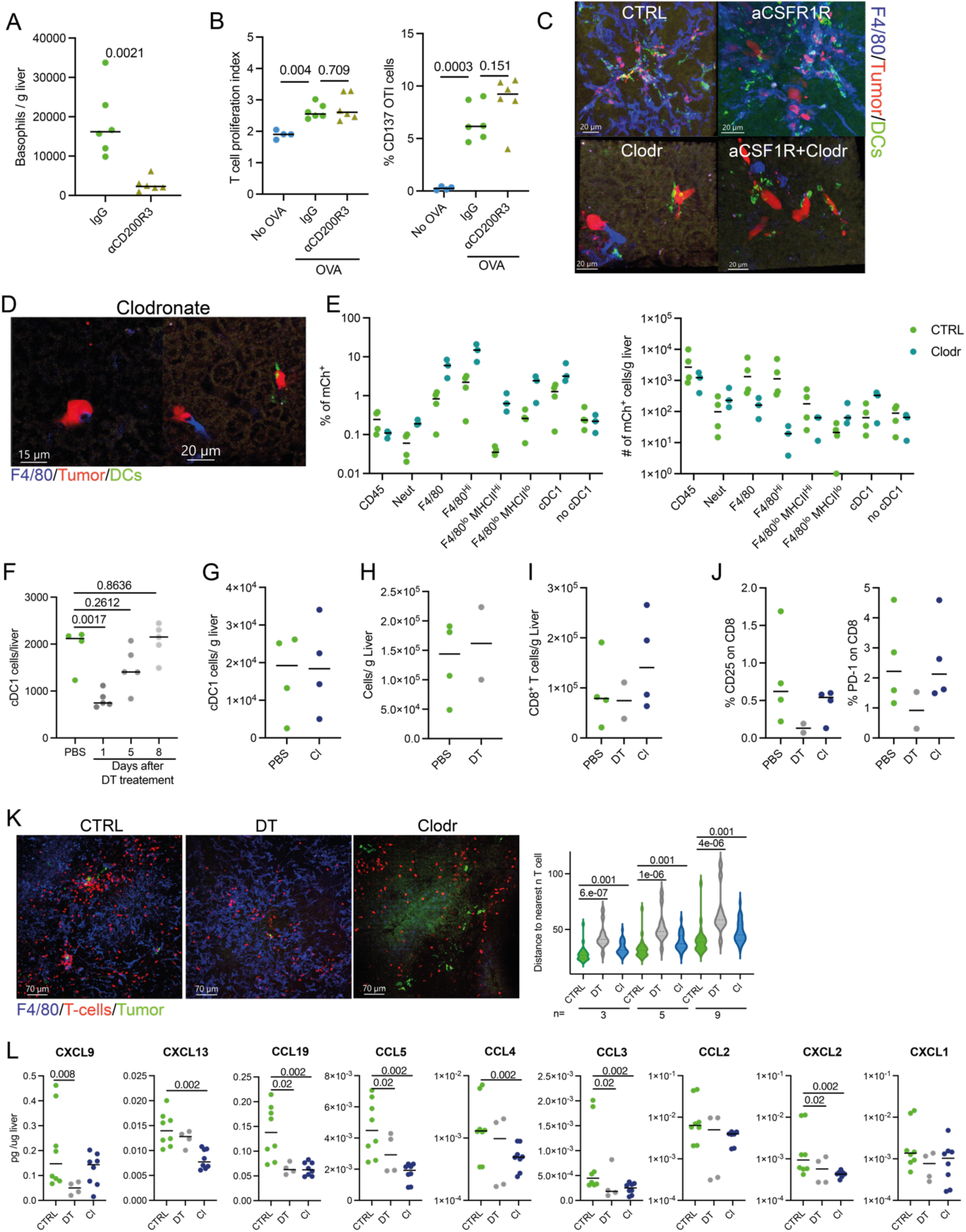
Immune cell requirements for efficient antigen cross-presentation of tumor metastatic cells in the liver. Related to Figure 6. In vivo cross-presentation assays with β2MKO CMT-OVA, as in Figure 5F, in mice pretreated with the CD200R3 basophil depleting antibody. (A) Assessment of depletion efficiency. (B) Flow cytometry assessment of adoptively transferred OTI cells’ proliferation (CTV dilution) and activation (CD137 surface expression). (C) Representative images of clarified CD11cEGFP livers stained for macrophages (F4/80) at the 24 hours post-metastatic time point in mice treated with different macrophage depletion regimens. (D) Representative images as in (C) but showing proper interaction of CD11c cells and F4/80^low^ macrophages with tumor cells in clodronate liposomes pretreated mice. (E) Flow cytometry quantification of mCherry^+^ myeloid cells upon clodronate depletion in the liver, shown as percentage within each population and absolute numbers within the liver. (F) Assessment of depletion efficiency of cDC1 in the liver over time in XCR1-DTR mice after a double DT treatment on days –2 and –1. (G) Flow cytometry assessment of cDC1s in the liver upon liver metastases elicitation in mice pre-depleted of macrophages with clodronate liposomes. (H) Flow cytometry assessment of macrophages in the liver upon liver metastases elicitation in XCR1-DTR mice pre-depleted of cDC1 cells. (I) Flow cytometry assessment of T cells in the liver upon liver metastases elicitation in cDC1 and macrophage-pre-depleted mice. (J) Flow cytometry assessment of T-cell activation markers CD25 and PD-1 in the liver upon liver metastases elicitation in cDC1 and macrophage-pre-depleted mice. (K) Representative images and T-cell-T-cell distance analysis of clarified livers from XCR1-DTRxCD2RFP 24h post-MC38 tumor metastasis. (L) Chemokine profile of the liver tissue 24h post-i.s. injection measured by multiplexed ELISA. Each dot represents a mouse, except for K, in which each dot represents a single measurement. In K, at least 5 FOV were analyzed per mouse. Medians are represented as a horizontal bar for each condition. Exact p-values are shown for statistical comparison, U Mann-Whitney tests, except for F, in which an ANOVA followed by a Dunnett’s post-hoc test was performed.

